# Single-cell DNA methylome and 3D genome atlas of the human subcutaneous adipose tissue

**DOI:** 10.1101/2024.11.02.621694

**Authors:** Zeyuan Johnson Chen, Sankha Subhra Das, Asha Kar, Seung Hyuk T. Lee, Kevin D. Abuhanna, Marcus Alvarez, Mihir G. Sukhatme, Kyla Z. Gelev, Matthew G. Heffel, Yi Zhang, Oren Avram, Elior Rahmani, Sriram Sankararaman, Sini Heinonen, Hilkka Peltoniemi, Eran Halperin, Kirsi H. Pietiläinen, Chongyuan Luo, Päivi Pajukanta

## Abstract

Human subcutaneous adipose tissue (SAT) contains a diverse array of cell-types; however, the epigenomic landscape among the SAT cell-types has remained elusive. Our integrative analysis of single-cell resolution DNA methylation and chromatin conformation profiles (snm3C-seq), coupled with matching RNA expression (snRNA-seq), systematically cataloged the epigenomic, 3D topology, and transcriptomic dynamics across the SAT cell-types. We discovered that the SAT CG methylation (mCG) landscape is characterized by pronounced hyper-methylation in myeloid cells and hypo-methylation in adipocytes and adipose stem and progenitor cells (ASPCs), driving nearly half of the 705,063 detected differentially methylated regions (DMRs). In addition to the enriched cell-type-specific transcription factor binding motifs, we identified *TET1* and *DNMT3A* as plausible candidates for regulating cell-type level mCG profiles. Furthermore, we observed that global mCG profiles closely correspond to SAT lineage, which is also reflected in cell-type-specific chromosome compartmentalization. Adipocytes, in particular, display significantly more short-range chromosomal interactions, facilitating the formation of complex local 3D genomic structures that regulate downstream transcriptomic activity, including those associated with adipogenesis. Finally, we discovered that variants in cell-type level DMRs and A compartments significantly predict and are enriched for variance explained in abdominal obesity. Together, our multimodal study characterizes human SAT epigenomic landscape at the cell-type resolution and links partitioned polygenic risk of abdominal obesity to SAT epigenome.

## Main

The global prevalence of abdominal obesity, defined as an excessive accumulation of adipose tissue in the abdominal region, has been increasing at an alarming rate over the past few decades^1,2^. Abdominal obesity is a known predictor of all-cause mortality, likely due to its increased risk of cardiometabolic disease (CMD), cardiovascular diseases, musculoskeletal diseases, certain types of cancers, and other adverse pathological conditions^3^. This has stimulated research interest in investigating the molecular origin of abdominal obesity and related co-morbidities by focusing on the subcutaneous adipose tissue (SAT), the key fat depot in expanding and buffering against obesity.

SAT is highly heterogeneous and comprises an array of cell-types^4^. Single nucleus RNA-sequencing (snRNA-seq) enables the discovery of cell-type level gene expression patterns in SAT^5,6^. However, this modality is limited to gene expression even though SAT function is also influenced by epigenomic processes, such as cytosine DNA methylation at CpG sites (mCG)^7^, and chromatin conformation^8^. Previous studies in other tissues have shown that cell-type level dynamic mCG in gene regulatory regions and gene bodies affect the expression of genes^9^. Furthermore, gene regulatory mechanisms need proper chromatin conformation, which is organized into compartments, domains, and loops^10^. However, cell-type level epigenomic landscape underlying the extensive heterogeneity in SAT is poorly understood in humans, which also hinders genetic risk assessment of abdominal obesity, the functional basis of which likely includes specific cell-type level epigenomic sites.

Single-nucleus methyl-3C sequencing (snm3C-seq) has emerged as a powerful and innovative platform to study DNA methylation and chromatin conformation at the cell-type resolution^11^. Recent studies identified cell-type level epigenomic signatures in various complex tissues in human, such as oocytes^12^, prefrontal^11^ and frontal cortex^13,14^, and other diverse brain regions^9^. Using a similar approach, previous studies have also comprehensively assessed the epigenomes of mouse brain cell-types^15–17^. However, cell-type level epigenomic signatures in the human key fat depot, SAT, are completely unknown. To address this important biomedical knowledge gap, we determined cell-type level DNA methylation, chromatin conformation, and gene expression signatures in SAT, assessed the involvement of methylation pathway genes in SAT cell-type level dynamic methylation patterns, identified cell-type level hypo-methylated region -associated transcription factor (TF) binding motifs, and investigated the contribution of variants in SAT cell-type level epigenomic sites to abdominal obesity risk.

## Results

### Overview of the study design

Epigenomic landscape of SAT is unknown at the cell-type level. To address this knowledge gap, we used snm3C-seq and snRNA-seq technologies on nuclei isolated from SAT biopsies from individuals with obesity (see Methods) (Fig. 1a) to generate cell-type level DNA methylation, chromatin conformation, and gene expression profiles in SAT. After performing careful quality control (QC) in each modality, we verified the high concordance of cell-type annotations derived from mCG and interaction modality as well as between mCG and gene expression. We then conducted analysis of differentially methylated regions to find cell-type level differences in DNA methylation patterns in SAT (Fig. 1b). To elucidate chromatin conformation dynamics in SAT cell-types, we systematically searched for cell-type level patterns in terms of the global contact distance distribution, as well as 3D genome features at various resolution (i.e., compartments, domains, and loops) (Fig. 1c). We next utilized cell-type level SAT snRNA-seq data (Fig. 1d) to investigate whether methylation pathway genes contribute to the discovered differences in DNA methylation patterns in SAT cell-types and cluster with adipogenesis pathway genes (Fig. 1e). We also identified cell-type-specific TF binding motifs associated with hypo-methylated regions of SAT cell-types (Fig. 1f). Finally, to understand how these cell-type level epigenomic differences relate to the key cardiometabolic phenotypes relevant to SAT, we examined whether variants in cell-type level DMRs and compartments contribute significantly to the polygenic risk of obesity and related cardiometabolic traits (Fig. 1g).

**Figure 1.**
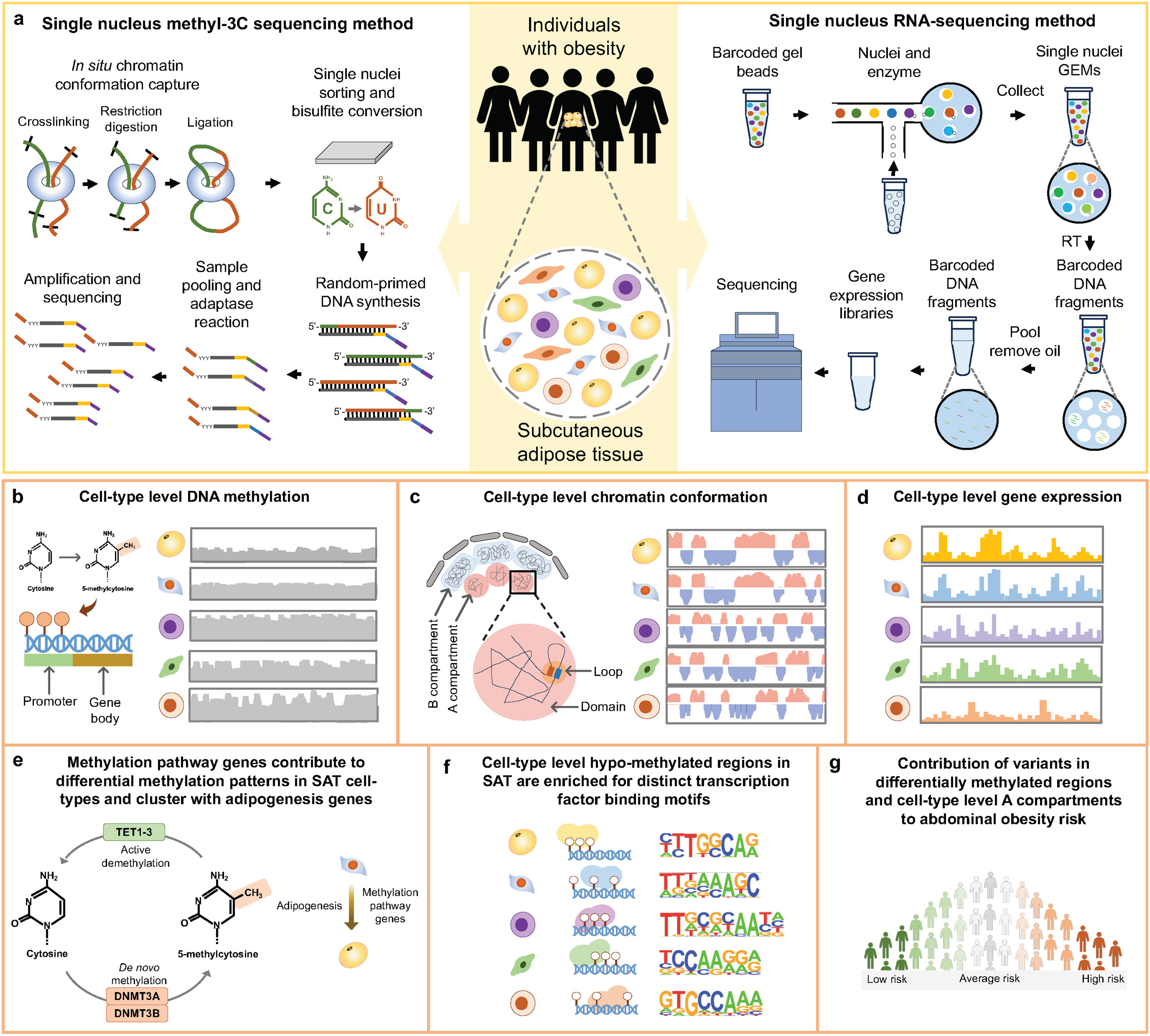
Schematic overview of the study design using single nucleus methyl-3C sequencing and single nucleus RNA-sequencing to profile cell-type level DNA methylation, chromatin conformation, and gene expression in the human subcutaneous adipose tissue (SAT) and partition the genetic risk of abdominal obesity. **a**, Illustration of single nucleus methyl-3C sequencing (snm3C-seq) and single nucleus RNA-sequencing (snRNA-seq) on nuclei isolated from SAT biopsies from females with obesity. **b-g**, Comprehensive analyses of DNA methylation, chromatin conformation, and gene expression profiles across the SAT cell-types to identify cell-type level differences in DNA methylation patterns (b) and chromatin conformation dynamics (c). Subsequently, we used the cell-type level SAT expression data (d) to determine whether methylation pathway genes contribute to the observed differences in methylation patterns in SAT cell-types and longitudinally cluster with adipogenesis pathway genes (e), identify cell-type-specific transcription factor (TF) binding motifs associated with hypo-methylated regions in SAT cell-types (f) as well as (e) to test the contribution of variants in cell-type level differentially methylated regions and A and B compartments to the genetic risk of abdominal obesity (g).

### Multimodal profiling of the SAT cells reveals highly concordant, yet partly asynchronous cell-type annotations among modalities

We used snm3C-seq to simultaneously profile single-cell level DNA methylation and chromatin conformation of nuclei isolated from five SAT biopsies (see Methods). A total of 6,652 nuclei passed our QC, with each cell having on average 2,215,680 non-clonal methylation reads and 236,850 chromatin contacts. We identified 7 main cell-types (adipocytes, adipose stem and progenitor cells (ASPCs), perivascular, endothelial, myeloid, lymphoid, and mast cells) using the global mCG of non-overlapping 5-kb bins and independently the intrachromosomal contacts among non-overlapping 100-kb bins (Fig. 2a). Interestingly, when analyzing the two modalities jointly to derive the *de novo* snm3C-seq annotation, we discovered a group of nuclei (n=63 nuclei), present in all 5 samples that demonstrated inconsistent cell-type annotations between the two modalities (i.e., categorized as perivascular cells by mCG and adipocytes by chromatin conformation) (Fig. 2b). We labeled them as the transitional cell-type cluster to highlight their potential developmental stage, observed using the two different omic profiles (Fig. 2a,b). During the differentiation of other tissues, the establishment of global chromatin 3D structure has previously been shown to precede the formation of methylation signatures^18^. In other words, the observed asynchrony between the mCG and conformation profiles suggests that the transitional cell-type cluster is undergoing active differentiation from perivascular cells to mature adipocytes, in line with recent studies that discovered perivascular adipocyte progenitors in mice and humans^19–21^.

**Figure 2.**
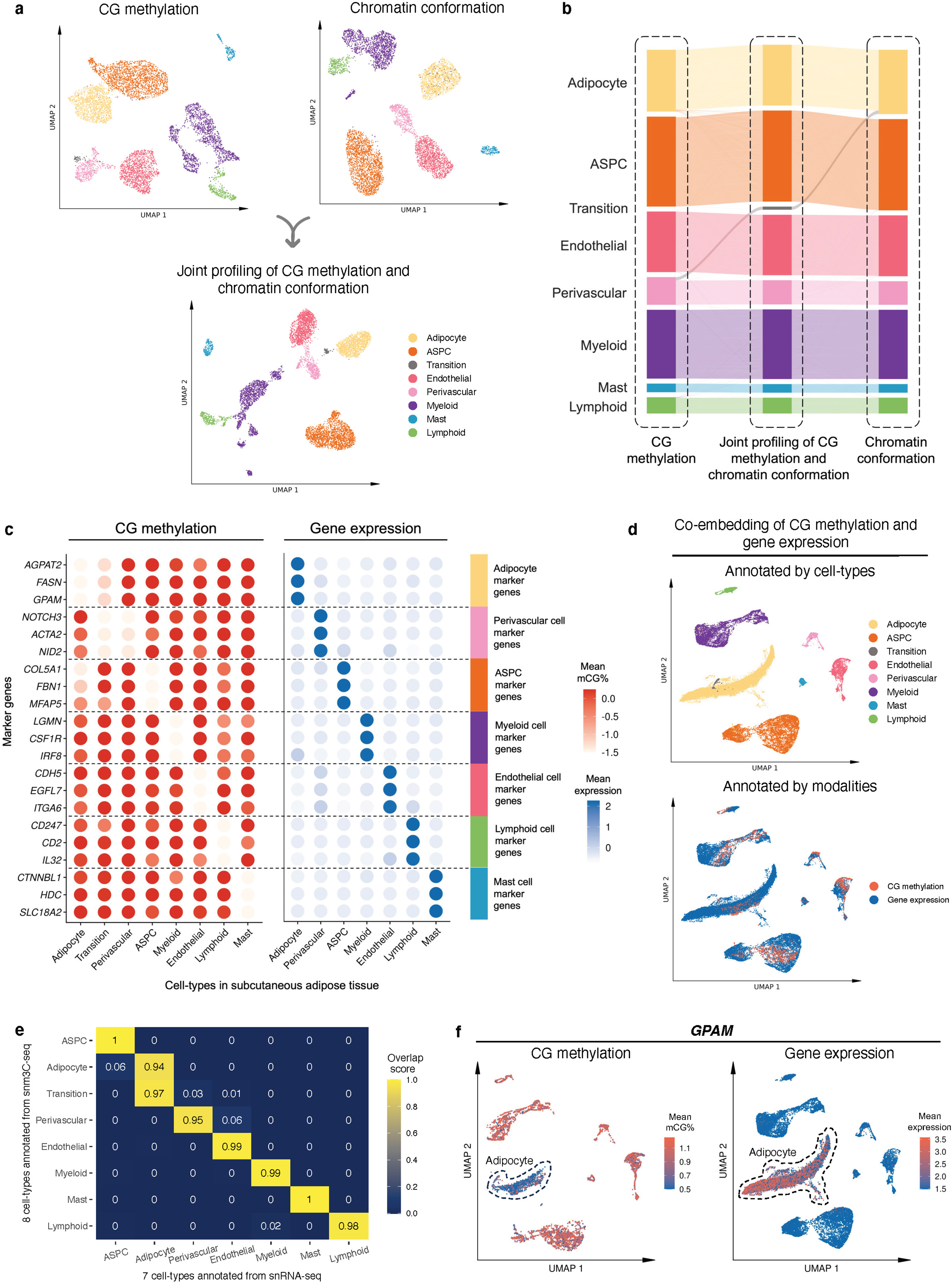
Single-nucleus level multi-omic profiles of SAT by jointly profiling methylation and chromatin conformation with snm3C-seq, followed by an integrative analysis with transcriptomic profiles, generated using SAT snRNA-seq. **a**, Dimension reduction of cells using 5-kb bin mCG (top left), 100-kb bin chromatin conformation (top right), and jointly integrating mCG and chromatin conformation (bottom), profiled by single nucleus methyl-3C sequencing (snm3C-seq) and visualized with uniform manifold approximation and projection (UMAP). Cells are colored by cell-types of subcutaneous adipose tissue (SAT). **b**, Sankey diagram showcases the high consistency among the SAT cell-type annotations derived from the 5-kb bin mCG (left), 100-kb bin chromatin conformation (right), and joint profiling of mCG and chromatin conformation (middle), with the exception of the transition cell-type cluster that is annotated as perivascular cells by mCG and adipocytes by chromatin conformation. **c-f**, Integrative analysis with snRNA-seq, evaluating the concordance of cell-type cluster annotations and cell-type marker genes across the used modalities. **c**, Comparison of gene-body mCG and gene expression profiles of cell-type marker genes across the matching SAT cell-types, independently identified within the respective modalities, excluding the expression profiles of the transition cell-type cluster that was not identified in the SAT snRNA-seq data. Dot colors represent the average gene-body mCG ratio normalized per cell (left), and the average log-transformed counts per million normalized gene expression (right). **d**, Co-embedding of snm3C-seq gene-body mCG and snRNA-seq gene expression, visualized with UMAP. Cells are colored by the SAT cell-types identified in **c** (top) and modalities (bottom). **e**, Concordance matrix comparing the snm3C-seq and snRNA-seq derived annotations, colored by the overlapping scores between the pairs of the SAT cell-types evaluated in the co-embedding space. **f**, UMAP visualization of the gene-body mCG ratio (left) and gene expression (right) for one adipocyte marker gene, *GPAM,* colored per cell similarly as in **c**. ASPC, adipose stem and progenitor cell.

We next investigated whether SAT snm3C-seq data can be integrated with SAT snRNA-seq data. First, we applied snRNA-seq on 29,423 SAT nuclei isolated from the same 5 SAT samples and 3 additional SAT samples from the Tilkka cohort (see Methods) to obtain the single-cell level expression profiles from matching SAT tissue, and annotated them at the cell-type resolution (Extended Data Fig. 1a). We then calculated the average gene-body mCG levels for all snm3C-seq nuclei as a proxy for their transcriptomic activity based on previous works that have found an inverse correlation between gene-body mCG level and expression level^14^. As expected, we observed strong and consistent correlations between gene-body mCG hypo-methylation and RNA expression across the identified cell-types (Fig. 2c). This correlation enabled us to integrate and co-embed the snm3C-and snRNA-seq cells, applying a mutual-nearest-neighbor based approach^22^ in the shared canonical component space (Fig. 2d, Extended Data Fig. 1b,c). Overall, these independently performed modality-specific annotations achieved a ≥0.94 overlap score across all cell-type pairs, in which a higher score indicates better integrated cells in the co-embedding space (see Methods) (Fig. 2e). Comparison between the snm3C-seq *de novo* annotation and its RNA-derived counterpart resulted in an adjusted rand index (ARI) of 0.975 and ≥0.95 confusion fraction (Extended Data Fig. 1d). Unique cell-type marker genes by these two modalities are shown in Supplementary Tables 1-2. Overall, the observed cell-type epigenome profiles, identified using the snm3C-seq, exhibit strong concordance with those derived from snRNA-seq transcriptome; however, at the same time they carry distinct modality-specific information. For example, the expression of a key adipocyte marker gene, *GPAM*, coincides with demethylation of the gene in the co-embedding space, which may allow for the recruitment of relevant proteins, e.g., TFs (Fig. 2f, Extended Data Fig. 2a-f). Moreover, the transition cell-type cluster was co-embedded close to the adipocytes profiled by the snRNA-seq (Fig. 2d,e, and Extended Data Fig. 1c,d), in contrast to its *de novo* global mCG annotation (i.e., perivascular cells) (Fig. 2b). Analysis of the gene-body mCG levels further revealed that it simultaneously shows diminished methylation levels on both adipocyte and perivascular marker genes (Fig. 2c, Extended Data Fig. 2g). In addition, when restricted to the relevant cell-types, the transition cell-type cluster could also be discerned from the low dimensional projection of the genome-wide 5-kb bin mCG profiles, i.e., in the absence of chromatin conformation information (Extended Data Fig. 2h). The above-mentioned mCG properties at the individual transcriptomics level and the global projection underscore the biological validity of the transition cell-type. In summary, mCG and chromatin conformation profiles generated by snm3C-seq robustly recapitulated epigenomic profiles of known major SAT cell-types, while also uncovering a subtle transition cluster, supporting the differentiation of human adipocytes also from the perivascular progenitors.

### Comparison of unique cell-type marker genes and their functional enrichments between gene-body mCG and gene expression modalities reveals both modality-specific and -shared molecular mechanisms

We first searched for differences in unique marker genes at the cell-type level between the gene-body mCG and gene expression modalities (Supplementary Tables 1-2) and found both modality-specific and -shared marker genes (Extended Data Fig. 3a). We observed that majority of the cell-type level marker genes were identified as modality-specific. For instance, 77 adipocyte marker genes are present in both modalities, while 286 are unique to gene-body mCG and 738 are unique to gene expression.

As the highly expressed cell-type marker genes can be involved in biological processes and pathways relevant to the cell-type function, we performed their functional enrichment analysis using the WebGestalt^23^ for mCG and gene expression modalities. We observed both shared and non-shared biological processes (Extended Data Fig. 3b) and KEGG pathways (Extended Data Fig. 3c) enriched among the adipocyte marker genes between mCG and gene expression modalities. However, although only 77 adipocyte marker genes (21% of mCG and 9% of gene expression markers) are present in both modalities, the majority of biological processes (63% of the pathways identified from mCG and 65% from gene expression) and KEGG pathways (65% of the pathways from both mCG and gene expression) are shared. For example, PPAR signaling pathway, a well-known adipose tissue pathway, is significantly (FDR<0.05) enriched among the adipocyte marker genes in both modalities. We also identified several other shared biological processes, including fat cell differentiation, enriched among the adipocyte marker genes. Enrichment of these shared biological processes and functional pathways between the two modalities suggests that both methylation and gene expression play roles in regulating cell-type-specific molecular mechanisms.

Next, we evaluated cell-type level methylation and gene expression of the PPAR signaling pathway genes (Fig. 3a) that are shared adipocyte marker genes between the two modalities. We observed that 6 genes of PPAR signaling pathway, *ACSL1*, *ADIPOQ*, *LPL*, *PCK1*, *PLIN1*, and *PLIN4*, are hypo-methylated in the adipocyte and transition cell-type, while the same genes are hyper-methylated in the rest of the cell-types in SAT. Our comparisons of the mean gene expression across SAT cell-types further revealed that these 6 genes of PPAR signaling pathway are predominantly expressed only in the adipocyte cell-type, with minimal expression in other SAT cell-types (Fig. 3a). Similarly, we observed that the fat cell differentiation genes, *ADIPOQ*, *LPL*, *LEP*, *TCF7L2*, *AKT2*, and *SREBF1,* are hypo-methylated and predominantly expressed in adipocytes compared to the other SAT cell-types (Extended Data Fig. 3d). These findings suggest that key genes of PPAR signaling pathway and fat cell differentiation are regulated by both transcriptional and epigenetic mechanisms.

**Figure 3.**
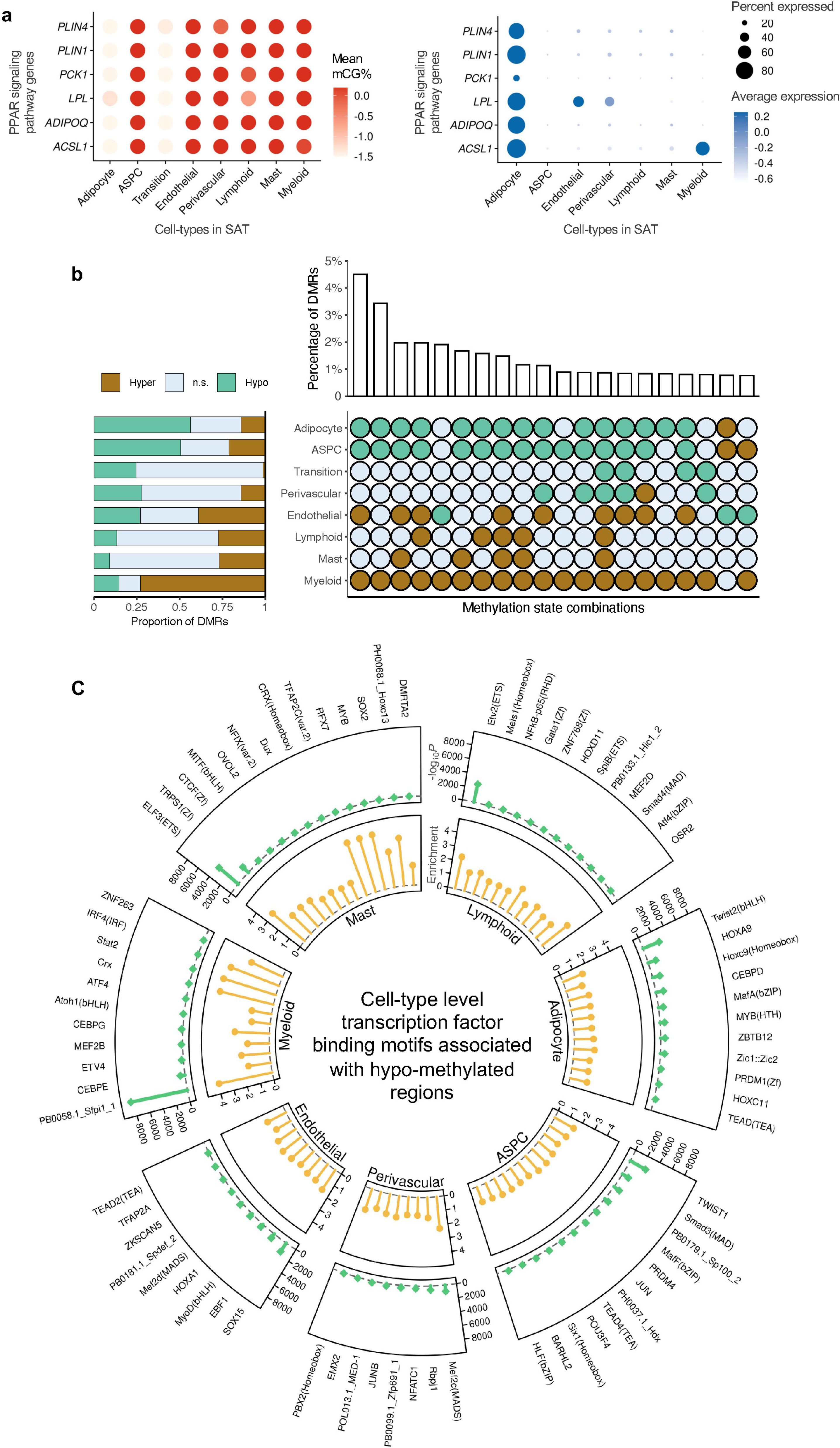
Functional pathways and gene regulatory potential of cell-type level gene-body mCG markers and differentially methylated regions. **a**, Dot plots of PPAR signaling pathway genes (*ACSL1*, *ADIPOQ*, *LPL*, *PCK1*, *PLIN1*, and *PLIN4*) that are shared adipocyte marker genes between the gene-body mCG and gene expression modalities, showing their gene-body mCG (left) and gene expression profiles (right) across the SAT cell-types. The color of the dot represents the mean percentage of mCG (left, red is high) and average expression of genes (right, blue is high), while the size of the dot represents the percentage of cells where the gene is expressed (right). **b**, Horizontal stacked bar plot (left) showing the marginal proportions of assigned methylation states across differentially methylated regions (DMRs) for each SAT cell-type (n.s. denotes non-significant) and upset plot (right) showing the top 20 combinations of methylation states across DMRs in decreasing order with their corresponding percentages. **c**, Circular plot summarizing the cell-type-specific transcription factor (TF) binding motifs associated with hypo-methylated regions in SAT cell-types. The outermost layer shows the names of cell-type-specific and significantly (*P*<1×10^-12^) enriched TFs in each SAT main cell-type. Track 1 shows the negative logarithmic of the *P* value (green lollipop) and track 2 shows the enrichment score (yellow lollipop). ASPC, adipose stem and progenitor cell, and FDR, false discovery rate.

### Analyses of the DMRs reveal striking differences in the number and abundance of hypo- and hyper-methylated regions between adipocytes and myeloid cells

To delineate the patterns of cell-type level DNA methylation in SAT, we identified genome-wide DMRs in 8 SAT cell-types (adipocytes, ASPCs, transition, perivascular, endothelial, myeloid, lymphoid, and mast cells) using methylpy^24,25^ (see Methods). Overall, 15.4% of the CG sites are differentially methylated across the SAT cell-types with a total of 705,063 CG DMRs covering 5.39% of the genome. These DMRs have a mean length of 220bp (SD=152bp) and consist of an average of 4.5 differentially methylated sites (DMSs) (SD=5.5). The large numbers of DMRs we identified in SAT cell-types support distinct cell-type level methylation patterns.

We observed striking genome-wide differences in the number and abundance of hypo- and hypermethylated regions among the SAT cell-types (Fig. 3b, Supplementary Table 3). In particular, of the total DMRs, 56.3% (n=396,758; -log_10_*P*=129 using one-tailed t-test, see Methods) and 50.6% (n=356,844; -log_10_*P* =104) are hypo-methylated in adipocytes and ASPCs, contrasting with only 14.6% (n=102,756) in myeloid cells. Conversely, we observed that up to 73.0% of the DMRs demonstrate hyper-methylation pattern in the myeloid cells (n=514,434; -log_10_*P*=163) versus merely 14.2% in adipocytes and 21.4% in ASPCs. Jointly investigating the differential methylation states across all cell-types revealed that 47.3% of the DMRs exhibit opposing profiles between adipocytes, ASPCs, and those of the myeloid cells. Taken together, our finding suggests that the widespread repression of regulatory activity in the myeloid cells is typically associated with heightened regulatory activity in adipocytes and ASPCs.

### Cell-type level hypo-methylated regions in SAT are enriched for distinct transcription factor binding motifs

To investigate the relevance of cell-type level hypo-methylated regions in gene regulation, we performed TF binding motif enrichment analysis using cell-type level hypo-DMRs. We first identified significantly (*P*<1×10^-12^) enriched TF binding motifs for each SAT cell-type using HOMER^26^. Hypo-methylated region -associated TFs and their corresponding enrichment ratios and *P* values are listed in Supplementary Table 4. Next, we searched for cell-type-specific TFs present in one cell-type and absent in others (Fig. 3c, Extended Data Fig. 4). Among the cell-type-specific TFs, the hypo-methylated regions in adipocytes are enriched for Twist family basic helix-loop-helix type transcription factor 2 (Twist2), homeobox A9 (HOXA9), and CCAAT enhancer binding protein delta (CEBPD); ASPCs for Twist family basic helix-loop-helix type transcription factor 1 (TWIST1), SMAD family member 3 (Smad3), and Jun proto-oncogene (JUN); and myeloid cells for CCAAT enhancer binding protein epsilon (CEBPE), Activating transcription factor 4 (ATF4), and Interferon regulatory factor 4 (IRF4). Our findings suggest that these cell-type-specific TFs might either bind to the DNA in a cell-type-specific manner or regulate cell-type level differential methylation patterns.

### Contact distance analysis identifies enrichment of short-range interactions in SAT adipocytes

Tight packaging of DNA inside the nucleus leads to physical contacts between genomic regions, which affects the gene expression machinery^27^. We observed substantial differences in the distribution of closely and distantly located interaction contacts at various genomic distances across the cells profiled by snm3C-seq (see Methods). In pairwise comparisons with other cell-types, adipocytes harbor significantly higher proportions of short-range interactions (100kb to 2Mb) compared to long-range interactions (10Mb to 100Mb), with the exception of the transition cell-type (–log_10_*P*>91; one-tailed Wilcoxon rank-sum test) (Fig. 4a,b). Specifically, the median proportion of the short-range interactions in adipocytes and the transition cell-type is 36.5%, whereas the median for others is 29.9%. Similarly, for long-range interactions, the median proportion is 26.0% both in adipocytes and the transitional cell-type, compared to 32.2% in others. This observation is in line with all cell-level clustering analyses using chromatin conformation information at various resolutions, which indicate that the transition cell-type shares more similarity with adipocytes (Fig. 2b, Fig. 4c,d, and Extended Data Fig. 5a,b).

**Figure 4.**
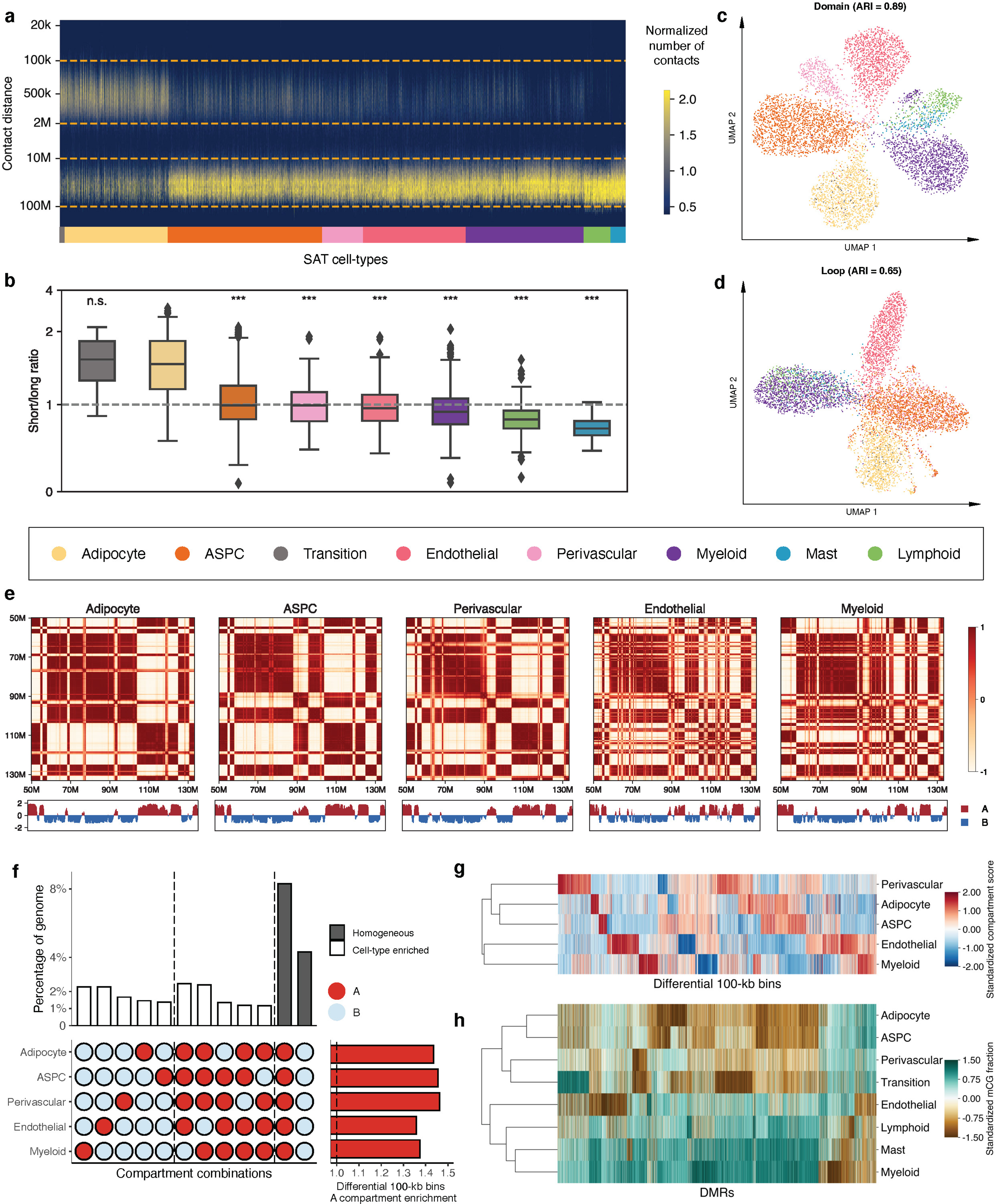
Analysis of chromatin conformation profiles in subcutaneous adipose tissue (SAT) reveals cell-type level diversity in compartments, domains, and loops. **a**, Frequency of contacts per cell against genomic distance. Cells are grouped by SAT cell-types and ordered by the median short to long-range interaction ratios. **b**, Short to long-range interaction ratios of SAT cell-types, ordered in the same way as in (a). Asterisks indicate the level of statistical significance of pairwise paired Wilcoxon test against adipocytes, and ***indicates -log_1*0*_*P*>50 and n.s. denotes non-significant. **c-d**, Uniform manifold approximation and projection (UMAP) visualization of low dimensional embeddings of cells using domains (c) and loops (d) as features, colored by the snm3C-seq annotation; adjusted rand index (ARI) evaluates the clustering concordance against snm3C-seq annotation. **e**, Heatmap visualization of the normalized interaction contact map on chromosome 12 and its corresponding compartment scores across SAT cell-types. **f**, Upset plot (left) visualizing a subset of the differential 100-kb bins (i.e., cell-type-specific and homogeneous compartment combinations) and their corresponding percentages; horizontal stacked bar plot (right) showing the marginal A compartment enrichment of differential 100-kb bins, stratified by cell-types. **g**, Dendrogram of the 5 most abundant SAT cell-types constructed with compartment scores on differential 100-kb bins. **h**, Similar to **g**, except on all annotated SAT cell-types, constructed with mCG fractions across DMRs. ASPC indicates adipose stem and progenitor cell.

In addition, the observed differences in the ratio of short to long-range interactions between ASPCs and adipocytes (Wilcoxon rank-sum –log_10_*P>*197) could suggest a link between the contact distances and functionally important genomic regions in adipogenesis. Given that ASPCs develop to adipocytes, we speculate that this change may reflect the unilocular lipid droplet formation in adipocytes that makes them larger than ASPCs.

### Global chromosomal conformation dynamics reflects lineage among SAT cell-types

Chromatin compartments, which connect stretches of the genome that are tens of mega-bases apart, reflect how cells arrange their chromosomal structures in three-dimensional space at the highest level^28^. We started our investigation of aggregated SAT cell-type level genome spatial topology by calculating the compartment scores on the pseudobulk contact matrices for the 5 most abundant cell-types (adipocytes, ASPCs, endothelial, perivascular, and myeloid cells) at 100-kb resolution. Based on the sign of the compartment scores, we partitioned the genome into either the active A compartment regions or the more repressive B compartment regions. The correlation matrices derived from the normalized interaction contact maps revealed visually distinct cell-type-specific plaid patterns. For example, on chromosome 6 (Extended Data Fig. 5c) and chromosome 12 (Fig. 4e), endothelial and myeloid cells harbor more intricate structures, indicated by the frequent compartment switches, whereas adipocytes, ASPCs, and perivascular cells tend to have longer stretches of region being annotated as the same compartment. Upon a closer inspection, a total of 11,571 100-kb bins, spanning 44.3% of the genome, are statistically differentially conformed among the 5 cell-types at an FDR<0.1 cutoff. The empirical FDR is estimated to be ≤0.02 (See Methods).

In each cell-type, differential 100-kb bins demonstrate at least 1.36-fold enrichment landing in the active A compartments compared to the genome-wide background (Fig. 4f, Extended Data Fig. 5d, and Supplementary Table 5). Interestingly, across all investigated SAT cell-types, the leading two predominant compartment combinations are the homogeneous A and B. These combinations account for 18.8% and 9.7% of the total differentially conformed regions (8.3% and 4.3% of the genome), suggesting significant heterogeneity within each compartment stratification (i.e., A and B). These are followed by combinations driven by myeloid and endothelial cells, either through cell-type-specific compartment flips (A->B or B->A) or coordinated flips involving both cell-types (Extended Data Fig. 5d). When focusing on compartment flips relative to adipocytes, a marked higher proportions of differential 100-kb bins categorized as adipocyte B compartment correspond to the A compartment of endothelial and myeloid cells (41.6% and 41.4%), in contrast to those of APSC and perivascular cells (28.0% and 33.8%, respectively; Extended Data Fig. 5e,f). Given that we observed similar distinct mCG patterns for endothelial and myeloid cells, we aimed to confirm whether the lineage dendrogram constructed from differential 100-kb bins would mirror the developmental trajectory inferred from DMRs. Indeed, hierarchical clustering consistently grouped endothelial and myeloid cells, characterized by pronounced hypo-methylation and frequent compartmental switches, into a distinct branch in both modalities (Fig. 4g,h), in line with a previous report showing that myeloid progenitors also give rise to vascular endothelial cells^29^.

### Cell-type specificity in regional 3D genome structures

In addition to compartmentalization, the genome maintains its finer spatial structure by forming interaction domains and cohesion-mediated chromatin loops. Analyzing snapshots of the 3D genome at 25-kb and 10-kb resolution allowed us to delineate these regional features at both cell and aggregated cell-type resolution. Besides the transition cell-type, adipocytes showcase an accumulation of significantly denser interaction domains (an average of 4,120 per cell, compared to 3,573 in others; -log_10_*P*>45, pairwise one-tailed Wilcoxon rank-sum test), while spanning a much shorter distance (median of 679,121bp per cell, compared to 783,213bp in others; Extended Data Fig. 6a-c). Interestingly, the number of detected domains is highly correlated with the ratio of short to long-range interactions (Pearson correlation coefficient=0.76; Extended Data Fig. 6d). This observation reinforces the idea that regional contacts are necessary to support the more intricate local 3D structures. Both features correlate with the general transcriptomic activity in the matching snRNA-seq data, where adipocytes show a 1.5-fold increase in the total UMIs (Extended Data Fig. 6e,f). Overlapping cell-type pseudobulk insulation scores with boundary probability, calculated as the fraction of cells with a boundary detected in a given cell-type, we identified a total of 1,791 differential boundaries (See Methods). Regarding chromatin loops, we detected a median of 47,837 and 5,797 cell-type level loop pixels and merged loop summits, respectively.

Adipocytes demonstrate a similar trend of having more loop summits (n=8,852) and a relatively shorter loop length (230,000bp, compared to others 290,000bp; Extended Data Fig. 6g,h). Along with the clustering results derived from regional interaction features (e.g., insulation scores, domains, and loops), which show highly concordant annotations (Fig. 4 c,d and Extended Data Fig. 5a,b), we conclude that granular 3D genomic features also exhibit significant heterogeneity across SAT cell-types.

### Influence of 3D topology on epigenetic regulation and associated gene-regulatory landscapes

The 3D topology of a cell also influences its transcriptomic dynamics with cell-type specificity. As expected, genes expressed in a cell tend to localize in its active A compartment, exhibiting ≥2.23 folds enrichment relative to the B compartment across the 5 most abundant cell-types. These ratios increase when restricted to the set of cell-type level unique marker genes (Supplementary Tables 1-2). Perivascular cells, in particular, exhibit a staggering 13-fold A/B ratio, leading to 92.8% of the unique marker genes landing in the A compartment (Supplementary Table 6). We next focused on ASPCs, a cell-type which undergoes active differentiation into adipocytes, and systematically evaluated how compartment flipping affects the downstream expression. On a global scale, 23.4% of the differentially conformed regions change from the ASPC A compartments to adipocyte B, while 27.5% convert from the B compartment to A. Residing within these topologically interesting regions are some key cell-type marker genes crucial for adipogenesis. For example, *COL1A2* and *LAMA2* are highly expressed in ASPCs and essential to the early stage of adipogenesis^30^, responsible for the extracellular matrix formation^31,32^. As the adipocytes mature, the regions harboring them flip to the B compartments, repressing the transcriptomic activity and halting the cell proliferation and tissue remodeling. On the other hand, the well-known adipocyte marker gene and adipokine, *ADIPOQ,* is located in an interaction domain unique to adipocytes with a pronounced demethylation pattern around its gene-body, likely facilitating additional TFs to bind and activate it functionally. The adjacent differential boundary marks a stretch of the genome, encapsulating 1Mb upstream and downstream of *ADIPOQ,* that transitions from the inactive B compartment in ASPCs to the active A compartment in adipocytes. Other strong *de novo* adipocyte marker genes, including *TENM3*, *CSMD1*, and *PCDH9*, also land within differential domains specific to adipocytes; additionally, *CSMD1* and *PCDH9* have cell-type-specific loop domains near the transcription starting sites (TSS). Together, our findings suggest that chromosome conformation, ranging from mega-base compartmentalization to kilo-base loop formation, reflects a higher level of carefully balanced coordination among the SAT cell-types, influencing both their epigenetic regulation profiles and trickling down to their transcriptional activities.

### DNA methylation pathway genes show cell-type preference in SAT expression, likely contributing to hyper- and hypo-methylation patterns in SAT cell-types

DNA methylation involves the covalent addition of a methyl group to DNA, a process facilitated by DNA methyltransferase enzymes, such as DNA methyltransferase 3 alpha (DNMT3A) and DNA methyltransferase 3 beta (DNMT3B)^33^. Conversely, DNA demethylation comprises the removal of this methyl group from the DNA by the ten-eleven translocation (TET) family proteins, specifically, TET1, TET2, and TET3 (Fig. 5a). To investigate whether methylation pathway genes regulate the observed difference in hyper- and hypo-methylation across the SAT cell-types, we analyzed the expression of DNA methylation and demethylation-related genes. Among the demethylase genes, *TET1* is preferentially expressed in adipocytes (-log_10_*P*>300; Wilcoxon rank-sum test) (Fig. 5b), in line with our observation that adipocytes have significantly more hypo-methylated (56.3%) than hyper-methylated regions (14.2%) (Fig. 5c,d). Among the DNA methyltransferases (DNMTs), our cell-type level SAT snRNA-seq data show that *DNMT3A* is predominantly expressed in myeloid cell-type with minimal or no expression in adipocytes (-log_10_*P*=131.9) (Fig. 5b). Consistent with our *DNMT3A* expression results, 73.0% of DMRs are hyper-methylated in the myeloid cells while only 14.2% are hyper-methylated in adipocytes (Fig. 5c). In addition, the expression of a methylation maintenance gene, *DNMT1,* is significantly lower in adipocytes than other SAT cell-types (-log_10_*P*=165.3) (Extended Data Fig. 7a). Expression of other methylation (*DNMT3B* and *UHRF1*) and demethylation genes (*TET2, TET3,* and *TDG*), shown in Extended Data Fig. 7a,b, suggest that *TET1* and *DNMT3A* are the most important genes that contribute to the observed adipocyte hypo-methylation and myeloid hyper-methylation patterns, indicating their potential mechanistic role in cell-type level DNA methylation signatures in SAT.

**Figure 5.**
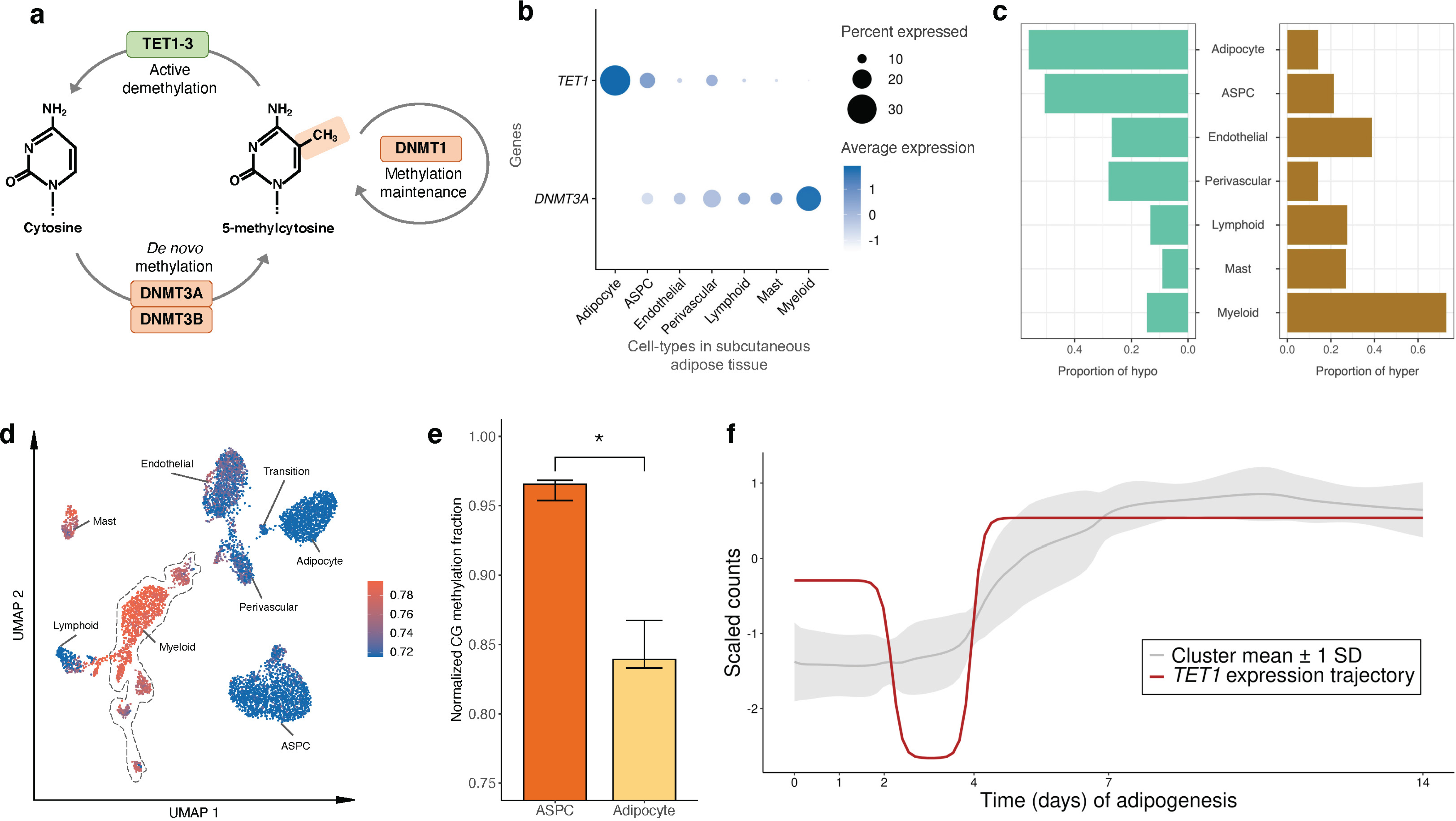
Analysis of mean gene expression and differentially methylated regions (DMRs) across subcutaneous adipose tissue (SAT) cell-types reveals the potential involvement of DNA methylation pathway genes in regulating cell-type level hyper- and hypo-methylation in SAT. **a**, A schematic representation of basic mechanisms and key players in DNA methylation and demethylation. **b**, Dot plot of *TET1* and *DNMT3A* showing their expression profiles across the SAT cell-types. The size of the dot represents the percentage of cells in which a gene is expressed within a cell-type and the color represents the average expression of each gene across all cells within a cell-type (blue indicates higher expression). **c**, Proportions of assigned hypo- (left) and hyper-methylated states (right) across DMRs. **d**, Uniform manifold approximation and projection (UMAP) visualization of the average global mCG ratio in a cell. **e**, Bar plot reflecting the distribution of normalized mCG fraction across genes that co-cluster with *TET1* in (f) for ASPCs and adipocytes. Asterisk indicates the level of statistical significance, *p≤0.05 using a paired Wilcoxon test. **f**, Longitudinal expression of *TET1* is plotted across the 14-day SAT preadipocyte differentiation. The ribbon behind the trajectory of *TET1* reflects the mean and standard deviation of the genes that clustered into similar trajectory patterns as *TET1* using DPGP. ASPC indicates adipose stem and progenitor cells.

### A demethylase gene, *TET1*, is temporally co-expressed with known adipogenesis genes across human primary preadipocyte differentiation

As temporal expression and co-expression patterns across human adipogenesis may relate to differential methylation between ASPCs and adipocytes, we examined longitudinal expression of 124 known adipogenesis pathway genes along with 5 demethylase and methylase genes (*UHRF1*, *TET1*, *TET2*, *TET3*, and *TDG*) across 6 time points of differentiation of human SAT primary preadipocytes (i.e., adipogenesis) (see Methods). We first observed that as expected, 121 of the tested known adipogenesis genes were longitudinally differentially expressed (DE) during SAT differentiation (adjusted *P*<0.05). To assess how the temporal co-expression patterns during adipogenesis relate to the expression of these demethylases and methylases, we clustered these genes using DPGP^34^ into 14 distinct clusters of longitudinally co-expressed genes (Supplementary Table 7). Notably, *TET1*, which we showed to be preferentially expressed in adipocytes when compared to the other methylase and demethylase genes (Fig. 5b and Extended Data Fig. 7a,b), clustered with known adipogenesis TFs and functionally important SAT genes, including *ADIPOQ*, *PLIN1*, *CEBPA*, and *LPL*. All exhibit significant demethylation and increased expression towards the end of adipogenesis (Fig. 5e,f). This suggests that *TET1* may function as a potentially important demethylation regulator of genes involved in adipogenesis.

### Variants in cell-type level DMRs and compartments are enriched for abdominal obesity risk

To understand the role of the identified differential epigenomic patterns in key cardiometabolic traits relevant to SAT, we explored the variants residing in the cell-type level DMRs and compartments for genetic evidence of contributions to cardiometabolic disease risk. Accordingly, we first examined whether the cell-type level DMRs contribute significantly to the polygenic risks of obesity traits, C-reactive protein (CRP), and metabolic dysfunction-associated steatotic liver disease (MASLD) by building annotated polygenic risk scores (PRS) for abdominal obesity (using waist-to-hip ratio adjusted for BMI (WHRadjBMI) as its well-established proxy^35,36^), BMI, MASLD, and CRP in the UK Biobank (UKB) cohort from variants landing in the cell-type level DMRs (see Methods). We observed that 6 of the WHRadjBMI PRSs, created from variants residing in adipocytes, ASPCs, and endothelial hypo-methylated, and endothelial, myeloid, and perivascular hyper-methylated DMRs, respectively, were not only significant predictors (p_R2_<0.05) of WHRadjBMI but also had significantly better incremental variance explained for WHRadjBMI (p_perm_10,000<0.05), compared to 10,000 permutated PRSs, each built with randomly selected clumped and thresholded SNPs with the same size as the DMR PRS (Fig. 6a). We additionally found that the CRP PRSs constructed from variants in three DMRs, including the myeloid hypo-methylated DMRs, were significantly enriched predictors for CRP (p_perm_10,000<0.05), while no enrichments were observed for BMI and MASLD.

**Figure 6.**
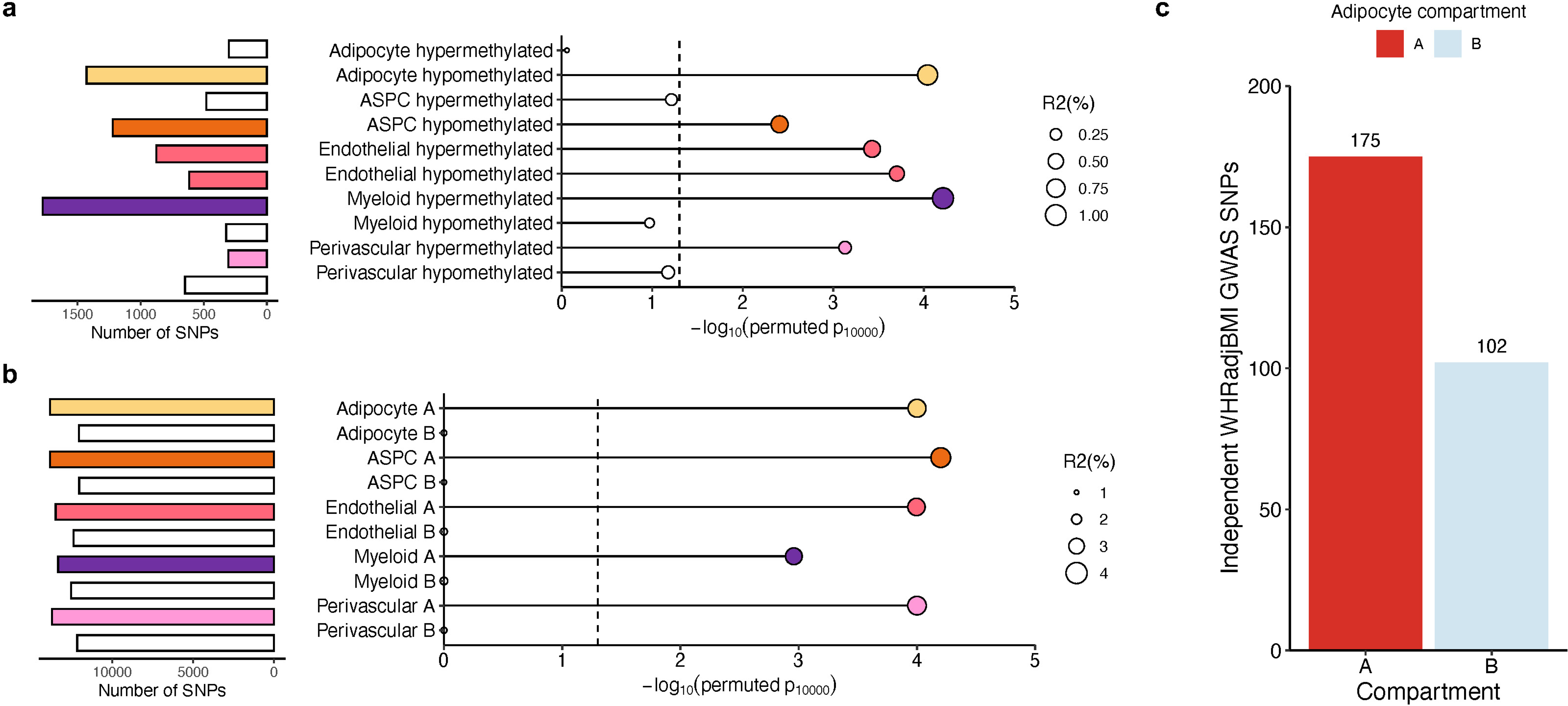
Partitioned abdominal obesity PRSs of several cell-type level DMRs and all cell-type level A compartments are enriched for variance explained in abdominal obesity, and 63.2% of non-redundant abdominal obesity GWAS variants land in adipocyte A compartment. **a-b**, Lollipop plots depict the incremental variance explained of each cell-type level PRS for abdominal obesity (using waist-hip-ratio adjusted for body mass index (WHRadjBMI) as a proxy) from the (a) DMRs, and (b) A and B compartments. Each lollipop represents a WHRadjBMI PRS, where the dot size corresponds to the incremental variance explained of the PRS. The grey vertical dotted line indicates the cutoff for significant enrichment of incremental variance explained (*P*_perm_10,000<0.05). On the left, horizontal bar-plots depict the number of SNPs used for the PRS construction. We color each bar and lollipop by the cell-type, where PRSs without a significant enriched PRS are outlined in grey without a filling. **c**, Bar plot showing the number of independent (r^2^<0.1) WHRadjBMI GWAS variants, passing genome-wide significance (*P*<5×10^-8^), from the WHRadjBMI GWAS, conducted in 195,863 individuals from the UK Biobank, grouped by the adipocyte compartment assignment. We shade each bar by compartment, where the A compartment is colored red and B compartment blue.

We then similarly studied the cell-type level compartments for enrichment of genetic risk. Due to the strong enrichments that we detected for WHRadjBMI among the cell-type DMR PRSs (Fig. 6a), we only constructed compartment-stratified PRSs for WHRadjBMI. For all five cell-types assessed, we noted that the PRSs built from variants residing in the A compartments were all consistently highly enriched predictors (p_perm_10,000<0.05) of WHRadjBMI, explaining ≥ 80% of the variance captured by the full genome (Fig. 6b; Supplementary Table 8). Conversely, we observed no such enrichment from the B compartment PRSs, suggesting the risk of abdominal obesity to be mainly driven by the A compartments. These abdominal obesity A compartment PRS results are further supported by the fact that 63.2% of the non-redundant abdominal obesity GWAS variants land in adipocyte A compartment (Fig. 6c and Extended Data Fig. 8a,b). Overall, our PRS results highlight the SAT cell-type level methylation and spatial conformation profiles as important contexts underlying the abdominal obesity risk.

## Discussion

Delineating cell-type level epigenomic landscape in human SAT is crucial for understanding their regulatory mechanisms and impact on obesity risk. We jointly profiled DNA methylation and 3D genome structure at single-cell resolution in SAT biopsies, which identified 705,063 DMRs with enriched TF binding motifs and cell-type level differential compartments. Our data revealed a highly dynamic reciprocal interplay between the SAT cell-type level epigenomes, particularly between adipocytes and myeloid cells. We further integrated the differential epigenomic sites with variant level data in the UK Biobank, thus uncovering their significant contributions to the polygenic risk of abdominal obesity. Finally, by integrating the cell-type level epigenomes with matching snRNA transcriptomes, we elucidated the potential role of specific methylation and demethylation pathway genes in the cell-type level differential methylation of human SAT.

The dynamic and asynchronous nature of SAT cell-types across modalities is exemplified by the identification of the transition cell population. Current evidence from perivascular adipose tissue in both mice and humans indicates that the perivascular adipocyte progenitor cells undergo adipocyte differentiation via induction of a thermogenic gene program^19^. Studies on rodent models has shown that *Ebf2*, a TF gene we also found to be hypo-methylated in the transition cell-type, is selectively expressed in mouse precursor cells of brown or beige fat^37^ and regulates thermogenic gene programming in mouse adipocytes^19,38^. This suggests that the observed transition cell-type represents the brown fat progenitor cells, which further differentiates into adipocytes in response to selective epigenomic and likely also environmentally driven changes.

Focusing on bulk methylome profiles, previous studies have reported differential DNA methylation patterns at the tissue level in SAT and their association with obesity^39,40^. However, underlying non-captured cell-type level methylation patterns and composition often confound tissue-level analyses^41^. Our SAT cell-type level methylation profiles and DMRs could serve as reference panels and provide informative features for computationally decomposing the heterogenous SAT mixtures^42–44^, a critical step for reducing false discoveries in tissue-level studies and facilitating cell-type-specific biomarker identification^42,45,46^. The cell-type composition itself could also hold significant clinical implications. For example, previous research using transcriptome profiling of perigonadal adipose tissue in mice and immunohistochemistry of human SAT has suggested that the accumulation of myeloid cells, particularly macrophages, correlates with increased adiposity^47^.

In our TF binding motif enrichment analysis, we observed cell-type-specific TF binding motifs that are enriched for hypo-methylated regions in SAT cell-types. Among the adipocyte-specific TFs, TWIST2 was identified as the top hit. A recent mice study reported that Twist2, a basic helix-loop-helix (bHLH) type TF, plays an essential role in lipid uptake and adipogenesis^48^. We also found an ASPC-specific TF, SMAD3, which acts as a downstream transcriptional transducer in the activin signaling pathway^49^. This pathway is well-studied for its role in the proliferation, differentiation, and function of preadipocytes^49,50^. Myeloid-specific TFs, CEBPE and ATF4, are known to regulate the expression of myeloid-specific genes^51^. These results endorse the possibility that both cell-type level hypo-methylation and TFs enriched in these hypo-methylated regions contribute to regulation of gene expression in SAT in a cell-type-specific manner.

Among the identified cell-type level 3D genome structures in SAT, adipocytes showcase a distinct regional topology, with a 1.51-fold enrichment in relative short-range interactions, 1.15-fold increase in the number of domains, and 1.72-fold increase in overall transcriptomic activity. Similar patterns have been observed in other human solid tissues and notably, these types of differences in the non-neuronal cells in brain have been linked to larger nuclear size^9,52,53^. Across all SAT cell-types, the widespread differences are reflected in the observed differential conformations detected with 44.3% of the compartment bins and 1,791 domain boundaries. The presence of the key adipocyte marker gene and adipokine, *ADIPOQ,* in a genomic region differentially conformed between ASPCs and adipocytes while heavily demethylated in adipocytes, further supports the idea that epigenomic structures reorganize during cell differentiation^54^, ultimately regulating downstream, regional, and cell-type level gene expression.

In our cell-type level investigations of methylation pathway genes, we found notably high expression of *TET1* in adipocytes and *DNMT3A* in myeloid cells, supporting a tissue and cell-type level reciprocal coordination and cross talk between the *TET1* expression and hypo-methylation in adipocytes and *DNMT3A* expression and hyper-methylation in myeloid cells. A previous study showed that *TET1* is an important DNA demethylase in adipose bulk tissue^55^. Another earlier study reported the involvement of *TET1* in adipocytokine promoter hypo-methylation in adipocytes^56^. Furthermore, previous studies have also demonstrated that both *TET1* and *DNMT3A* compete to regulate epigenetic mechanisms^57^. Thus, our findings are in line with these previous results, and taken together with our new cell-type level results endorse the possibility that *TET1* and *DNMT3A* play a crucial role in regulating the dynamic epigenetic landscape across the SAT cell-types.

Abdominal obesity is highly polygenic^58^. Previous studies have successfully built predictive genome-wide PRSs for abdominal obesity^59,60^ and shown that a high genetic predisposition to abdominal obesity predicts regain of abdominal obesity following weight loss^58^. However, less is known about the characteristics of specific genomic regions that contribute most to the polygenic risk of abdominal obesity, which could ultimately improve individual disease risk assessment. By constructing the partitioned PRS scores of abdominal obesity based on the two epigenomic single-cell level modalities, we discovered that variants in both adipocyte DMRs and A compartments significantly predict abdominal obesity and are enriched for variance explained in abdominal obesity using 10,000 permutations. This indicates that epigenomic sites in SAT adipocytes harbor significant polygenic risk for abdominal obesity.

Our study has some limitations. First, as the study comprises Finnish females with obesity, inclusion of males and individuals with normal weight would help elucidate potential sex-specific epigenomic landscapes and differences across the various BMI categories. Second, larger and more diverse set of samples could provide insight into population-based differences in the underlying epigenomic complexities in SAT cell-types. Third, inclusion of visceral adipose tissue (VAT) data into future studies would uncover cell-type level methylation and chromatin conformation patterns in this metabolically important other adipose depot as well as their differences when compared to SAT. However, requiring a surgical and medically indicated procedure, VAT biopsies are more invasive, thus making them practically less feasible. Nevertheless, taken together our study provides a valuable insight into the cell-type level epigenomes in human SAT to be followed up in future studies.

## Methods

### Tilkka cohort

Eight Finnish females with obesity underwent abdominal SAT liposuction at Tilkka Hospital, Helsinki, Finland. We performed snRNA-seq on all 8 SAT biopsies and snm3C-seq on 5 SAT biopsies. The study was approved by the Helsinki University Hospital Ethics Committee and all participants provided a written informed consent. All research conformed to the principles of the Declaration of Helsinki.

### UK Biobank cohort

For our genome-wide association study (GWAS) enrichment and polygenic risk score (PRS) analyses, we used genotype and phenotype data from the 391,701 unrelated individuals of European-origin of the UK Biobank cohort (UKB)^61,62^. As describes previously^61,62^, data for UKB were collected across 22 assessment centers. Genotype data were obtained using one of either the Applied Biosystems UK BiLEVE Axiom Array or Applied Biosystems UK Biobank Axiom Array, and imputed with the Haplotype Reference Consortium and the merged UK10K and 1000 Genomes phase 3 reference panels^61,62^. Data from UKB were accessed under application 33934.

### *In situ* chromatin conformation capture and fluorescence-activated nuclei sorting

We performed in situ chromatin conformation capture (3C) using an Arima Genomics Arima-HiC Kit as previously described^53^ with the following modification: the amount of Triton-X 100 in the NIBT buffer was increased to 1% to account for the large amount of fat in adipose tissue. The fluorescence-activated nuclei sorting (FANS) and library preparation were performed using the snmC-seq3 workflow (https://www.protocols.io/view/snm3c-seq3-kqdg3x6ezg25). The snmC-seq3 libraries of human SAT was sequenced using the Illumina NovaSeq 6000 instrument with S4 flow cells generating 150 bp paired-end reads. The sequencing reads of snm3C-seq were mapped using Taurus-MH^11^ (https://github.com/luogenomics/Taurus-MH).

### Snm3C-seq quality control and preprocessing

We filtered the cells profiled by snm3C-seq based on the following metrics: 1) the estimated non-conversion rate mCCC%<0.015; 2) the global mCG%>0.5; 3) the global mCH%<0.15; 4) the total number of interaction contacts >100,000 and <500,000; and 5) at least one intra-chromosome contact present in each autosome after filtering out reads with either end mapped to the ENCODE blacklist region^63^.

### Genotype quality control and imputation in the Tilkka cohort

We genotyped the DNAs from the Tilkka participants using the Infinium Global Screening Array-24 v1 (Illumina). In our quality control (QC), we used PLINK v1.9^64^ to remove 1) individuals with missingness >2%, 2) unmapped, strand ambiguous, and monomorphic SNPs, and 3) variants with missingness >2% and Hardy-Weinberg Equilibrium (HWE) *P* value<10^-6^. In addition, we imputed biological sex using the ‘--sex-check’ function in PLINK v1.9^64^ and confirmed that they matched the reported sex for all individuals.

We utilized the HRC reference panel version r1.1 2016^65^ to perform genotype imputation against on the Michigan imputation server. Before imputation, we removed duplicate variants, as well as variants with allele mismatch with the HRC reference panel and matched strand flips or allele switches to match the panel before haplotype phasing using Eagle v2.4^66^. To perform the genotype imputation, we used minimac4^67^ and performed QC on the data by removing SNPs with imputation score R^2^<0.3 and HWE *P* value<10^-6^.

### Nuclei isolation and snRNA-seq of human SAT in the Tilkka cohort

We performed SAT snRNA-seq experiments on the snap-frozen SAT biopsies from the Tilkka participants, as previously described^59^. We measured the concentration and quality of nuclei, separately for each sample, using Countess II FL Automated Cell Counter after staining with trypan blue and Hoechst dyes. To construct the libraries, we used the Single Cell 3’ Reagent Kit v3.1 (10x Genomics) and analyzed the quality of cDNA and gene expression using Agilent Bioanalyzer. We sequenced the libraries from each participant together on an Illumina NovaSeq S4 with a target sequencing depth of 600 million read pairs.

To maximize the samples size of the snRNA-seq data in the Tilkka cohort, we performed the joint snRNA- and snATAC-seq experiment on the subset of 5 SAT biopsies and included the snRNA-seq data in this study. Briefly, we combined 300 mg of the 5 SAT biopsies into a gentleMACS C tube (Miltenyi Biotec) containing 3 ml of chilled 0.1X lysis, including 10 mM Tris-HCl, 10 mM NaCl, 3 mM MgCl2, 0.1% Tween-20, 0.1% IGEPAL CA-630, 0.01% Digitonin, 1% BSA, 1 mM DTT, and 1 U/μL RNase inhibitor. We next dissociated the tissues by placing the gentleMACS C tube on the gentleMACS Dissociator (Miltenyi Biotec) and running the ‘4C_nuclei_1’ program. The tissues were incubated in the lysis buffer for a total of 15 minutes including the time on the dissociator. After the incubation period, we added 3 ml of chilled wash buffer, containing 10 mM Tris-HCl, 10 mM NaCl, 3 mM MgCl2, 1% BSA, 0.1% Tween-20, 1 mM DTT, and 1 U/μL RNase inhibitor, to the lysate and filtered the lysate mixture through a 70 μm MACS strainer, followed by a 30 μm MACS strainer. Next, the nuclei were centrifuged at 300 rcf for 5 minutes at 4°C and the supernatant was removed without disrupting the nuclei pellets. We then resuspended the nuclei pellet in 3 ml of chilled wash buffer and passed through a 30 μm MACS strainer. The final concentration and quality of nuclei were measured using the Countess II FL Automated Cell Counter after staining with trypan blue and Hoechst dyes and the snRNA-seq library was constructed using the Single Cell Multiome ATAC + Gene Expression Reagent Kit (10x Genomics). We used the Agilent Bioanalyzer to assess the quality of cDNA and sequenced the library on an Illumina NovaSeq SP with a target sequencing depth of 400 million reads.

### Processing of the SAT snRNA-seq data from the Tilkka cohort

First, we aligned the raw snRNA-seq data from all experiments against the GRCh38 human genome reference and GENCODE v42^68^ annotations with STAR v2.7.10b^69^. We utilized the ‘--soloFeatures GeneFull’ option to account for full pre-mRNA transcripts. Then the quality of the raw and mapped snRNA-seq data were evaluated using FastQC. To remove empty droplets as well as nuclei with high levels of ambient RNA, we ran DIEM v2.4.0^70^ with initialization parameters 1) UMI cutoffs ranging from 100 to 1000 to define debris, and 2) *k=*50 for the initialization step with k-means clustering, along with all other default parameters. We applied the sample specific UMI cutoffs in the initialization step to account for differences in sequencing depth between samples. Next, we removed clusters with low average UMIs, low average number of unique genes detected (nFeatures), high percentage of mitochondrial mapped reads (%mito), and high number of mitochondrial and ribosomal genes as top expressed features. Droplets with nFeatures≤200, UMI≤500, %mito≥10, and spliced read fraction≥90% were removed using Seurat v4.3.0^71^. Next, we used Seurat v4.3.0^71^ to log-normalize gene counts employing the ‘NormalizeData’ function; identify top 2,000 variable genes using the ‘FindVariableFeatures’ function; scale the gene counts to mean 0 and unit variance using the ‘ScaleData’ function; perform principal component analysis (PCA) using the ‘RunPCA’ function; and cluster the nuclei with a standard Louvain algorithm, using parameters of the first 30 PCs, and a resolution of 0.5, respectively.

To remove reads from ambient RNA molecules, we ran DecontX^72^ with the removed low-quality nuclei as the background and the Seurat cluster assignment as the ‘z’. We then removed nuclei with nFeatures≤200, UMI≤500, UMI≥30,000, and %mito≥10 based on the remaining reads. For the multiplexed snRNA-seq data of 5 SAT biopsies from the joint snRNA-and snATAC-seq experiment, we ran demuxlet from the popscle software tool^73^ to identify the originating individual of each nucleus. Next, DoubletFinder^74^ was employed to remove predicted doublets. Since DoubletFinder requires a predicted number of doublets as input, we used a pN-pK parameter sweep, as previously recommended^74^, to select pN=0.25 and the most optimal pK value that maximizes the mean-variant normalized coefficient.

### Snm3C-seq data integration, clustering, and annotation

While methylation features could be stratified into mCH and mCG, mCH is primarily found only in the human brain and not elsewhere in the body (e.g., SAT)^75,76^. Thus, we represented only the mCG profiles of each cell by 5-kb bins across autosomal chromosomes. Briefly, per cell and for each 5-kb bin, we calculated a hypo-methylation score (i.e., the *P* value of observing fewer methylated reads under a binomial distribution with the expected probability of a methylated read set to the global mCG rate of the cell, and the number of trials set to the coverage of the 5-kb bin). We next binarized the score matrix by converting nominally significant entries (i.e., *P* value<0.05) to 1 and the rest to 0, as described previously^16^. Bins that overlapped with the ENCODE blacklist region^63^ were excluded from the clustering analysis. Next, we performed latent semantic indexing (LSI) on the term-frequency, inverse-log-document-frequency transformed matrix, implemented in the ALLCools package^15^ (v.1.0.23) to obtain the mCG profile embedding, and then further omitted the first dimension due to its high correlation with sequencing depth. For the chromosome conformation modality, we imputed the contact matrix of each cell at 100-kb resolution using scHicluster^77^ (v.1.3.5) with pad=1 and used singular value decomposition (SVD) to project all intra-chromosome contacts between 100kb to 10Mb that land in autosomal chromosomes to a low dimensional space. To remove the sample level batch effect, we applied Harmony^78^ (v.0.0.9) on the snm3C-seq joint embedding (i.e., the concatenation of the top 10 dimensions from both modalities). The resulting matrix was used for k-NN graph construction (k=25), Leiden consensus clustering, and uniform manifold approximation and projection (UMAP) visualization.

We annotated the clusters *de novo* by leveraging the negative correlation between mCG and transcriptional activity^9,14,15^ and the known SAT marker genes, reported previously^79^. Specifically, we calculated the average mCG fractions of the gene-body and normalized the fractions per cell by first taking the posterior of the mCG probability of each gene with a Beta distribution prior, representing the genome-wide mCG rate of the cell, before scaling by the inverse of it^15^. Thus, hypo-methylated SAT marker genes, characterized by normalized scores notably lower than the genome-wide average of 1, suggest strong expression patterns.

We conducted modality-specific clustering, visualization, and annotation similarly, while calculating them using information solely from their respective embedding. All clusters were merged to the resolution representing the canonical major cell-types identified in SAT (adipocytes, ASPCs, perivascular, endothelial, mast, myeloid, and lymphoid cells), with the exception of the “transition” cluster, which was categorized as adipocytes if using only the chromosome conformation information and perivascular cells if using only the mCG profiles.

### SnRNA-seq data integration, clustering, and annotation

We integrated all remaining high-quality droplets from all snRNA-seq data with reciprocal principal components analysis (rPCA) implemented in Seurat v4.3.0^71^ and clustered integrated data with a standard Louvain algorithm, using parameters of the first 30 PCs, and a resolution of 0.5. We annotated each cluster with their cell-type using SingleR v1.8.1^80^ with a previously published single-cell atlas of human SAT as a reference^79^.

### Co-embedding of snm3C-seq and snRNA-seq data

We aligned the snm3c-seq cells as the query with snRNA-seq cells as the reference under the canonical correlation analysis (CCA) framework of Seurat v.4.1.0^22^, similarly as described previously^17,81^. To capture the shared variance between modalities, we started with reversing the sign of the normalized gene-body mCG fractions, and then applied CCA between the resulting matrix and the expression count matrix on the set of genes used to integrate the RNA datasets, while also requiring >5 mapped reads in snm3C-seq cells. Transfer anchors were identified within the top 30 canonical component space as the top 5 mutual nearest neighbors. We further filtered and weighted the anchors by distances in the snm3C-seq joint embedding to impute RNA-based annotations and expression profiles of the snm3C-seq cells. ARI was used to evaluate the concordance between the *de novo* annotation and the imputed RNA-based annotation. Cells profiled by both technologies were merged on their imputed expression profiles, projected to low dimensional space with PCA, and visualized by UMAP (constructed on the top 10 PCs).

For a given *de novo* snm3C-seq and the snRNA-seq annotated cell-type cluster pair, we defined the overlap score as the sum, across all clusters in the shared CCA co-embedding space, of the minimum proportion of cells in each modality-specific cluster that overlapped with a co-embedding cluster. Thus, the overlap score ranges from 0 to 1, where 0 indicates a complete separation and 1 indicates a perfect co-localization of modality-specific cells within the same co-embedding cluster. We normalized the multi-class confusion matrix per row by the number of cells to derive the confusion fractions. The confusion matrix was calculated by comparing the *de novo* annotations of the snm3C-seq cells with their intermediate imputed cell-type labels, which were determined through weighted votes from transfer anchors using snRNA-seq as reference.

### Cell-type level SAT marker gene identification in snm3C-seq and snRNA-seq

We excluded the following genes from differential testing of the cell-type marker genes: 1) genes that overlap with the ENCODE blacklisted regions; 2) smaller genes (≤200 bp) mostly covered by other genes (overlap region≥90% of the gene length)^15^, and 3) genes with a shallow coverage, constantly methylated or un-methylated, defined as those without ≥10 methylated or un-methylated counts in ≥10 cells belonging to the cell-type under investigation. Cell-type level differentially methylated genes (DMGs) were determined *de novo* by performing the Wilcoxon rank-sum test on the normalized gene-body mCG fractions of the snm3C-seq cells in a one-vs-rest way. We retained genes that had Benjamini-Hochberg (BH) adjusted *P* value<0.05, and at the same time exhibited a hypo-methylation difference of ≥0.1 in terms of the average normalized fraction when compared to the other cell-types^15^. For transcriptomics, we first filtered for the set of expressed genes in SAT, defined as those with ≥3 counts in ≥3 cells^72^. We used ‘FindAllMarkers’ function in Seurat to identify marker genes employing the default parameters with the exception that we constrained our search to the subset of positive marker genes with ≥25% non-zero expression in either the tested cell-type or the other ones^82–84^. Subsequently, we filtered out genes with BH-adjusted *P* values≥0.05. To obtain unique marker genes on both snm3C-seq and snRNA-seq, we removed genes identified as marker genes for more than one cell-type.

### Pathway enrichment analyses for SAT cell-type marker genes in gene-body mCG and gene expression modalities

To identify cell-type level biological processes and functional pathways enriched among the cell-type marker genes in mCG and gene expression modalities, we utilized the web-based tool WebGestalt^23^ that identifies the overrepresentation of gene sets in Gene Ontology (GO) biological processes and Kyoto Encyclopedia of Genes and Genomes (KEGG) pathways. For each SAT cell-type, we used the unique cell-type marker genes as the input, with only the genes expressed within that cell-type as the reference for the enrichment analysis. Biological processes and KEGG pathways with FDR<0.05 were considered statistically significant.

### Cell-type level methylation profile analysis

To obtain the cell-type level mCG profiles, we aggregated single-cell level number of CG methylated counts and total coverage based on the snm3C-seq joint annotation and further merged reads mapped to adjacent CpG in +/-strands. We then used MethylPy^24,25^, implemented in the ALLCools package, to detect genomic regions that display distinct mCG patterns across various cell-types. Differentially methylated sites (DMSs) on autosomes were tested across all 8 annotated cell-types using default parameters. For all DMSs, we assigned one of the three states per cell-type, hypo-, neutral-, or hyper-methylated, based on whether the fitted residual (i.e., the normalized deviation away from the mean methylation level) fell below the 0.4, between the 0.4 and 0.6, or above the 0.6 quantile of its chromosome-wide background, respectively^15^. Nearby DMSs (within 250bp) with Pearson correlations of >0.8 for the methylation fractions across the cell-types were merged into differentially methylated regions (DMRs). Differential methylation states were assigned to each DMR based on the average of those of the DMSs it encompasses. DMRs containing only one DMS or without any hyper-hypo-methylation state assignment, and DMRs or DMSs overlapping ENCODE blacklist regions were excluded from downstream analyses.

Additionally, we tested whether DMRs showcase genome-wide cell-type preferential differential methylation states under a regression framework. Specifically, for every differential state (hyper- or hypo-methylation), we fitted the following regression model across all cell-types and all autosomal chromosomes. The response variable, the cell-type level fraction of the DMRs with the desired methylation state on a chromosome, was modeled by several independent variables, including the log normalized number of cells belonging to the corresponding cell-type and a one-hot encoded indicator for all cell-types in SAT. This approach allowed us to quantify the cell-type level contribution to the methylation state fraction while also calibrating for the inherent fraction differences induced by the varying statistical power arising from the differences in the coverage among cell-types. The stratification to hyper- and hypo-methylation states naturally suggests a directionality in the test. Thus, we report the log10 *P* values derived from one-tailed t-tests, evaluating the probability of observing a larger positive contribution on the cell-type indicator variable under the null.

### Prediction of cell-type-specific transcription factor binding motif using HOMER

We performed TF binding motif enrichment analysis using the motif discovery tool HOMER v4.11.1 (Hypergeometric Optimization of Motif EnRichment)^26^. For each SAT main cell-type, we used the hypo-methylated regions as input data for motif enrichment analysis with the HOMER function ‘findMotifsGenome.pl’. TF binding motifs with *P*<1×10^-12^ were considered statistically significant. Circular visualization of cell-type-specific TF binding motif enrichment results was prepared using the circlize package^85^ in R.

### Cell-type level compartment analysis

Based on the snm3C-seq joint annotation, we merged scHicluster imputed single-cell level contact matrices at 100-kb resolution per chromosome to form the cell-type level pseudobulk conformation profiles for the 5 most abundant cell-types (adipocytes, ASPCs, endothelial, perivascular, and myeloid cells) as well as a cell-type aggregated version. For each chromosome independently, genomic bins in the cell-type aggregated contact map with abnormal coverage, defined as the total number of interactions between itself and all other bins, were removed from the compartment analysis. Specifically, we kept bins with a coverage <99th percentile and above twice the 50th percentile minus the 99th percentile. This filtration typically removes poorly mapped regions like telomere, centromere, and blacklisted regions^9^. Cell-type level pseudobulk conformation profiles were then normalized by the distance between the contacts and converted to correlation matrices by dcHic v2.1^54^. For all 5 cell-types, we fitted PCA on the resulting matrices per chromosome and extracted the first two PCs as candidates of the compartment scores. The dcHic tool heuristically selected the PC that maximized the absolute correlation with transcription start site (TSS) and CpG density as the compartment scores and, if, needed, flipped its sign to ensure that regions with positive scores corresponded to more active (A) compartments. We visually inspected the compartment scores to verify that they indeed captured the plaid pattern instead of the chromosome arms. Compartment scores from all 5 cell-types were then quantile normalized. Finally, we tested for genomic bins that demonstrated large deviations away from the cell-type average under a multivariate normal distribution, measured by the Mahalanobis distance using the covariance matrix learned with outlier bins removed. Bins with FDR corrected *P* values<0.1 were labeled as differentially conformed regions^54^. Empirically, we observed FDR<0.02 when repeating the same analysis but only on a null set of cell-type level contact maps, obtained by arbitrarily shuffling the annotation of the cells before merging to the pseudobulk level (i.e., a scenario where any differential compartment detected is false positive by construction), indicating a conservative calibration of the testing result by dcHic.

### Characterizing interaction domains and chromatin loops in SAT

For interaction domains, we used scHiCluster v.1.3.5^77^ with pad=2 to impute contact matrix of each cell per autosomal chromosome at a 25-kb resolution, restricted to contacts within 10Mb. We detected domains for each cell using TopDom^86^ and calculated the insulation scores across all 25-kb genomic bins with a window size of 10 bins using the imputed contact profiles. We then projected the domain boundaries with LSI and insulation scores with PCA into low-dimensional space. Similar to snm3C-seq joint annotation, we applied Harmony v.0.0.9^78^ to correct for batch effects and visualized the top 10 low-dimensional embeddings with UMAP. Cell-type level domain boundary probabilities were calculated as the fraction of cells with a detected domain boundary in a given 25-kb bin across all cells belonging to the specified cell-type. Differential domain boundaries were evaluated per bin based on criteria similar to those described previously^53^. Specifically, we required the Z-score transformed chi-square statistic >1.960 (97.5 percentile of standard normal distribution), the differences between the maximum and minimum cell-type boundary probabilities to be >0.05, detection as a local boundary peak (maximum), simultaneous detection as a local insulation score valley (minimum), and finally, FDR<0.001.

To analyze chromosomal looping, we used scHiCluster v.1.3.5^77^ with pad=2, window_size=30000000, and step_size=10000000 to impute contact matrix of each cell per autosomal chromosome at a 10-kb resolution, restricted to contacts within 10Mb. Loop pixels were detected from cell-type pseudobulk imputed contact profiles based on enrichment relative to both its global and local backgrounds. We aggregated near-by loop pixels passing an empirical FDR of 0.1 to loop summits. To create cell-level embeddings based on looping features, we first gathered all identified loop pixels and built a binary cell-by-loop matrix, where each entry indicates whether at least one contact was detected in the cell at the corresponding loop pixel^9^. We used LSI to project the cell-by-loop matrix, Harmony to correct for batch effect, and UMAP to visualize the top 10 low-dimensional embeddings.

### Human primary preadipocyte (PAd) differentiation experiment

We previously cultured cryopreserved human primary SAT preadipocytes (Zen-Bio catalog # SP-F-2, lot L120116E) for adipogenesis (14-day preadipocyte differentiation) and conducted ATAC-seq and RNA-seq across 6 time points: 0d, 1d, 2d, 4d, 7d, and 14d^6^. Briefly, for each time point and modality, we plated cells at confluency to create 4 isogenic replicates. Libraries for RNA-seq were prepared using the Illumina TruSeq Stranded mRNA kit and sequencing was performed on one lane of Illumina NovaSeq S1 flowcell. We obtained an average of 42M +/-5M (SD) reads per sample.

### Longitudinal differential expression (DE) across six human adipogenesis time points

We analyzed the longitudinal trajectory patterns of 124 known pathway genes involved in adipogenesis (https://www.wikipathways.org/pathways/WP236.html) and expressed in these adipogenesis data, as well as 5 additional demethylase and methylase genes (*UHRF1*, *TET1*, *TET2*, *TET3*, and *TDG*) also expressed in these data using ImpulseDE2 v0.99.10^87^ across the 6 adipogenesis time points. We used the runImpulseDE2 function with parameters *boolCaseCtrl*=FALSE, *boolIdentifyTransients*=TRUE, and *scaNProc*=1 on the respective RNA-seq gene expression counts. All *P* values were corrected for multiple testing using FDR<0.05.

### Identification of longitudinal trajectories of co-expressed adipogenesis genes and their methylation regulators

To search for longitudinal co-expression patterns among the key demethylase and methylase genes across human adipogenesis, we ran DPGP v0.1^34^ to cluster genes by their expression trajectories. We only included the genes that were identified as significantly longitudinally DE (FDR<0.05) during human adipogenesis from ImpulseDE2^87^, as described above.

### Construction of partitioned cardiometabolic polygenic risk scores for cell-type level DMRs and compartments

To assess the contributions of the SAT cell-type level differentially methylated regions (DMRs) and cell-type level compartments on the genetic risk for cardiometabolic traits, we constructed partitioned polygenic risk scores (PRSs) for each DMR for body mass index (BMI), waist-hip-ratio adjusted for BMI (WHRadjBMI), C-reactive protein (CRP), and metabolic dysfunction-associated steatotic liver disease (MASLD) in the UKB^61,62^, using the imputed MASLD status by Miao et. al^88^ for MASLD, and for WHRadjBMI for each cell-type compartment set. Only annotations from adipocytes, stromal, myeloid, endothelial, and perivascular cells were examined.

We first generated GWAS summary statistics for each trait with a 50% base group (n= 195,863) by applying a rank-based inverse normal transform to each trait and used the linear-mixed model approach of BOLT-LMM v2.3.6^89^, including age, age^2^, sex, the top 20 genetic PCs, testing center, and genotyping array as covariates. Variants with MAF<1% and INFO<0.8 were removed from the summary statistics. We then partitioned the remaining 50% into a 30% target and 20% validation groups for developing and applying the PRS model, respectively. Variants with MAF<1% and INFO<0.8 were removed from the used GWAS summary statistics, and the variants missing in >1% subjects, with MAF<1%, or violating Hardy Weinberg equilibrium as well as the individuals with >1% genotypes missing or extreme heterozygosity were removed from the target and validation genotype data^90^.

To compute the PRS for each outcome, we first generated independent marker sets by performing LD-clumping on all QC passing variants in the genome using plink^91^, with an LD R2 threshold of 0.2, and a window size of 250-kb. We then used the 30% test set (n=115,120) to identify the optimal *P* value cut point at the genome-wide level. Briefly, we applied the plink^91^ –score functionality to separately compute aggregated scores from subsets of the genome-wide clumped SNPs passing a range of a *P* value threshold from 5×10^-8^ to 0.5, using effect sizes and *P* values from the GWAS summary statistics. After identifying the best thresholding cutoff in the 30% test set (0.05 for WHRadjBMI and CRP, 0.3 for BMI, and 0.2 for MASLD), we computed regional PRSs in the 20% validation set (n=76,758), consisting of the clumped and thresholded SNPs landing within the DMR or compartment. Variance explained (R2) by each PRS, were calculated by adjusting each trait for age, age^2^, the top 20 genetic PCs, testing center, genotyping array, and sex, applying a rank-based inverse-normal transform, and then regressing the PRS on the adjusted trait.

To evaluate the significance of the variance explained by the PRS, we performed a permutation analysis, in which we randomly selected from the set of genome-wide clumped and thresholded SNPs, 10,000 sets of SNPs of the same size as the clumped and thresholded SNPs, overlapping with the DMR or compartment, and compared their R2 to the R2 of the 10,000 permutations.

## Data availability

The data that support the findings in this manuscript are available from the UK Biobank. However, restrictions apply to the availability of these data, which were used in this study under UK Biobank Application number 33934. UK Biobank data are available for bona fide researchers through the application process: https://www.ukbiobank.ac.uk/learn-more-about-uk-biobank/contact-us. The snm3C-seq and snRNA-seq data from the Tilkka cohort will be made available in the NIH Gene Expression Omnibus (GEO) upon acceptance, under accession number GSEXX. The bulk RNA-seq data from the primary human preadipocyte differentiation experiment was previously made available in GEO, under accession number GSE249195.

## Code availability

All packages and software used in this study were from their publicly available sources, as outlined in the Methods.

## Acknowledgements

We would like to thank the participants of the Tilkka cohort and the UK Biobank. This study was supported by NIH grants R01HL170604 (P. P.), R01DK132775 (P. P.), and R01HG010505 (E. H., P. P.). This research was conducted using the UK Biobank Resource under application number 33934.

## Author contributions

Z. J. C., S. S. D., and P. P. conceptualized and designed the project. Z. J. C., S. S. D., A. K., S. H. T. L., K. D. A., M. A., M. G. S., and K. Z. G. carried out the computational analyses. M. G. H., O. A., E. R., S. S., E. H., C. L., and P. P. suggested the analyses, provided support to perform them and participated in the discussion of results. M. A., Y. Z., S. H. T. L., C. L., and P. P. generated the snm3C-seq data, snRNA-seq and the bulk RNA-seq data. S. H., H. P., and K. H. P. collected the cohorts and samples. P. P. and E. H. funded the omics data generation and computational resources. Z. J. C., S. S. D., A. K., S. H. T. L., M. G. S., and P. P. wrote the manuscript. All authors read, reviewed, and/or edited the manuscript.

## Extended Data

**Extended Data Figure 1.**
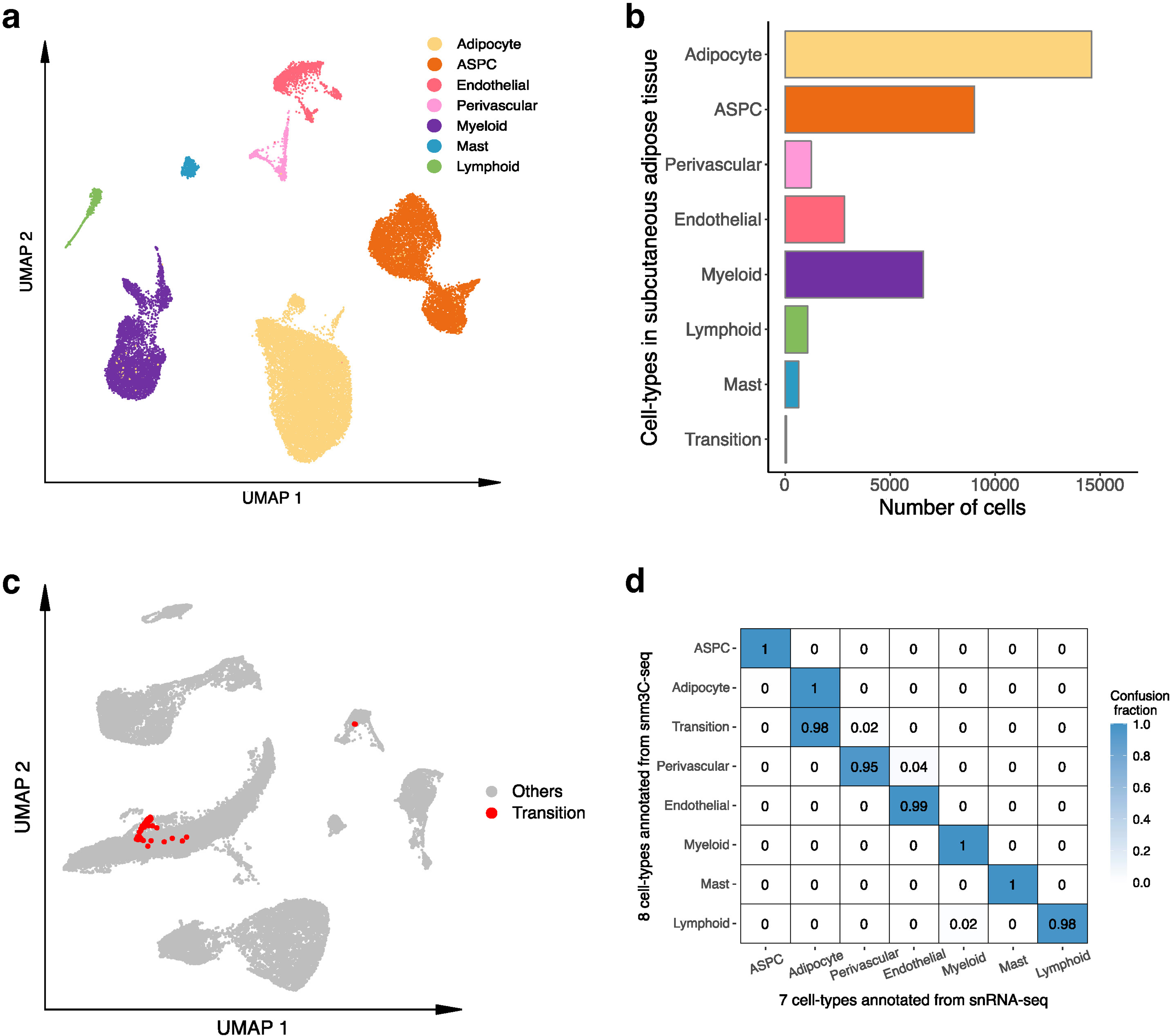
Integrative analysis between subcutaneous adipose tissue (SAT) cells profiled by single nucleus methyl-3C sequencing (snm3C-seq) and single nucleus RNA sequencing (snRNA-seq). **a**, Dimension reduction of cells (n=29,423) profiled by snRNA-seq and visualized with uniform manifold approximation and projection (UMAP). **b**, The total number of cells profiled by snm3C-seq and snRNA-seq stratified by the SAT cell-types. **c**, Co-embedding of snm3C-seq gene-body mCG and snRNA-seq gene expression, visualized with UMAP, highlighting the transition cell-type in red and other SAT cell-types in grey. **d**, Confusion matrix comparing the concordance between the *de novo* snm3C-seq annotations (row) and the snRNA-seq-derived annotations (column). The confusion fraction is calculated as the multi-class confusion matrix normalized by the cell counts per row. ASPC indicates adipose stem and progenitor cells.

**Extended Data Figure 2.**
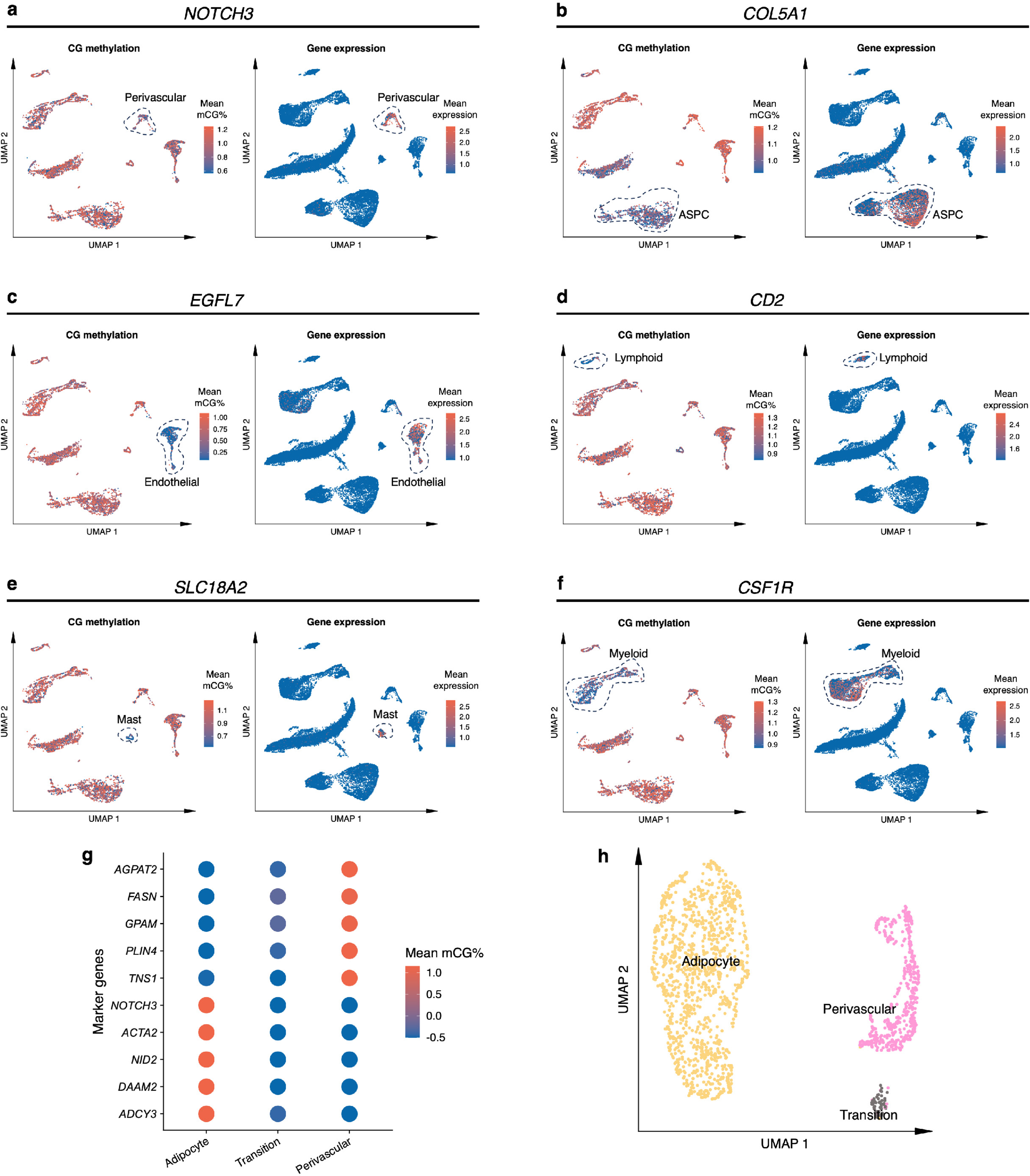
Gene-body mCG and RNA expression profiles across SAT cell-type marker genes and clustering analysis of the transition cell-type. **a-f**, Uniform manifold approximation and projection (UMAP) visualization of the gene-body mCG ratio, normalized per cell (left) and log-transformed counts per million normalized gene expression (right) for perivascular marker gene *NOTCH3* (a), ASPC marker gene *COL5A1* (b), endothelial cell marker gene *EGFL7* (c), lymphoid cell marker gene *CD2* (d), mast cell marker gene *SLC18A2* (e), and myeloid cell marker gene *CSF1R* (f). **g**, Gene-body hypo-methylation of adipocyte marker genes (top 5 rows) and perivascular cell marker genes (bottom 5 rows) across adipocytes, perivascular cells, and the transition cell-type. Dot colors represent the average gene-body mCG ratio normalized per cell. **h**, Dimension reduction of cells profiled by snm3C-seq and restricted to adipocytes, perivascular cells, and the transition cell-type, using exclusively the 5-kb bin mCG profiles and visualized with UMAP. ASPC indicates adipose stem and progenitor cells.

**Extended Data Figure 3.**
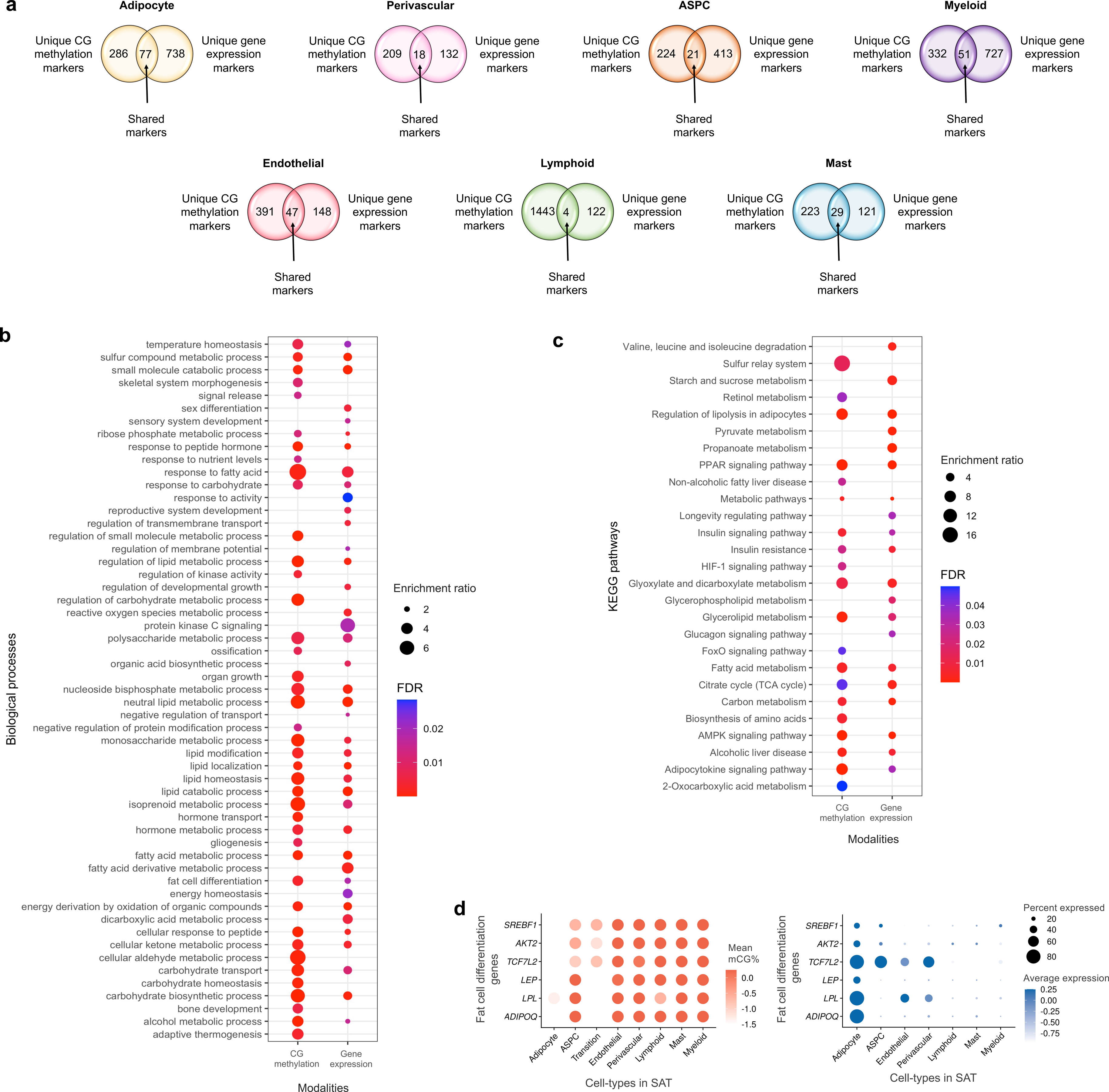
Comparisons of unique cell-type marker genes in SAT cell-types, and biological processes and functional pathways enriched among the adipocyte marker genes between the gene-body mCG and gene expression modalities. **a**, Venn diagrams showing the number of shared and modality-specific unique SAT cell-type marker genes (adipocytes, perivascular cells, ASPCs, myeloid cells, endothelial cells, lymphoid cells, and mast cells) between the gene-body mCG and gene expression modalities. **b-c**, Dot plots showing significantly (FDR<0.05) enriched biological processes (b) and KEGG functional pathways (c) using unique adipocyte marker genes in gene-body mCG and gene expression modalities. The size of the dot represents the enrichment ratio for biological processes (b) and KEGG functional pathways (c), while the color of the dot indicates FDR (blue is highly significant) (b-c). **d**, Dot plots of fat cell differentiation biological process genes (*ADIPOQ*, *LPL*, *LEP*, *TCF7L2*, *AKT2*, and *SREBF1*) that are shared adipocyte marker genes between the mCG and gene expression modalities, showing their gene-body mCG (left) and gene expression profiles (right) across the SAT cell-types. The color of the dot represents the mean percentage of mCG (left, red is high) and average expression of genes (right, blue is high), while the size of the dot represents the percentage of cells where the gene is expressed (right). ASPC indicates adipose stem and progenitor cells and FDR, false discovery rate.

**Extended Data Figure 4.**
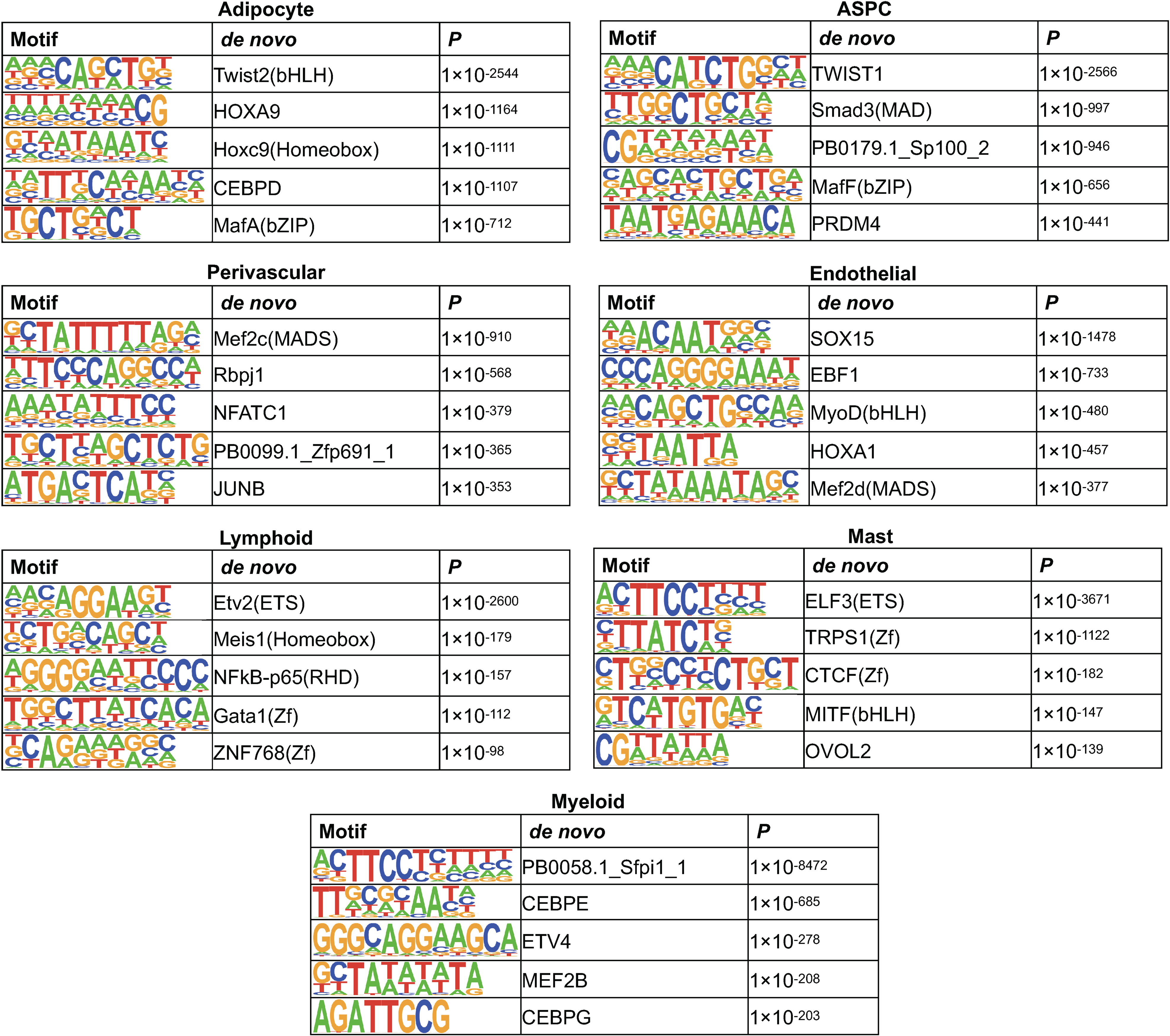
Cell-type level hypo-methylated regions are enriched for specific transcription factor (TF) binding motifs. We show the top five cell-type-specific TF binding motifs (sorted by *P*) that are enriched among the hypo-methylated regions of the SAT cell-types, identified using HOMER motif enrichment analysis. ASPC indicates adipose stem and progenitor cells.

**Extended Data Figure 5.**
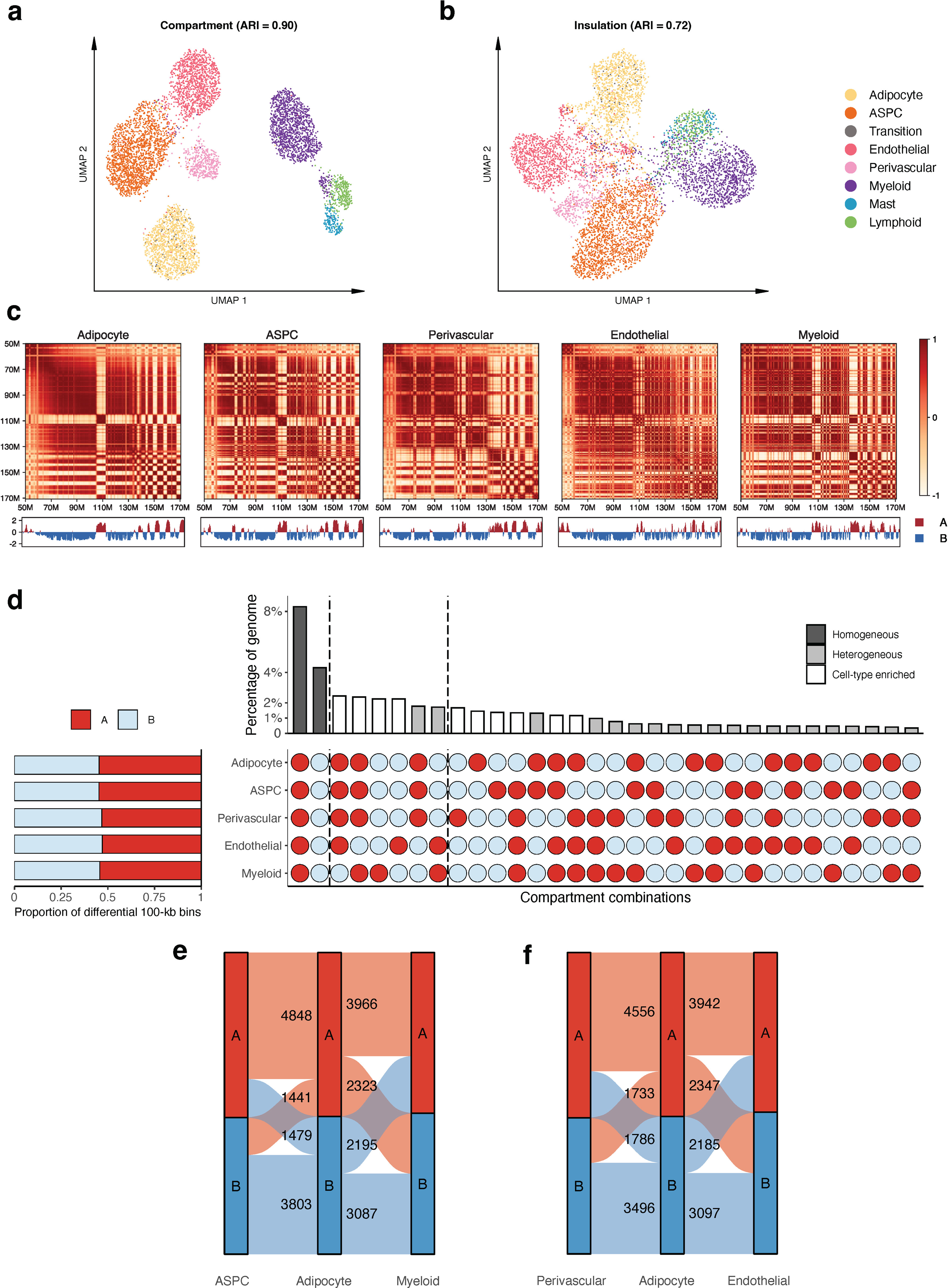
Cell-type level differences in chromatin conformation of subcutaneous adipose tissue (SAT). **a-b**, Uniform manifold approximation and projection (UMAP) visualization of low dimensional embeddings of cells using compartment (a) and insulation scores (d) as features, colored by the snm3C-seq annotation. Adjusted rand index (ARI) evaluates the clustering concordance against snm3C-seq annotation. **c**, Heatmap visualization of the normalized interaction contact map on chromosome 6 and its corresponding compartment scores across the SAT cell-types. **d**, Horizontal stacked bar plot (left) showing the marginal proportions of differential 100-kb bins stratified by their annotated A and B compartments in the 5 most abundant SAT cell-types and upset plot (right) showing all compartment combinations across differential 100-kb bins in decreasing order with their corresponding percentages (Homogeneous, Cell-type enriched, and Heterogeneous correspond to unique A or B compartment in 0, 1, or more than 1 cell-types, respectively). **e**, Sankey diagram breaking down of the numbers of differential 100-kb bins annotated as A (red) and B (blue) compartment belonging to ASPCs (left), adipocytes (middle), and myeloid cells (right). **f**, Similar to **e**, except on perivascular cells (left), adipocytes (middle), and endothelial cells (right). ASPC indicates adipose stem and progenitor cells.

**Extended Data Figure 6.**
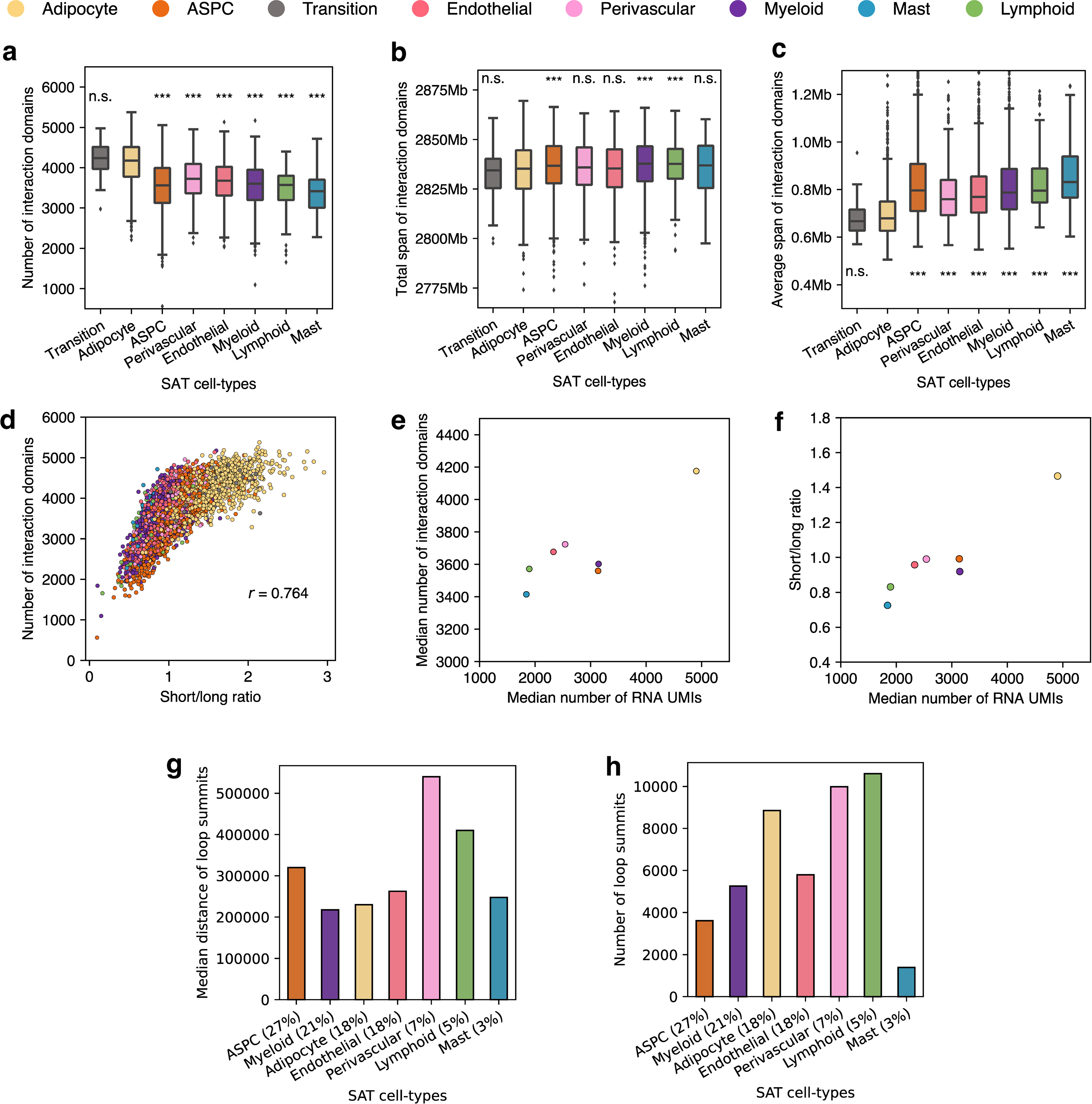
Cell-type specificity in interaction domains and loops. **a-c**, Box plots visualizing the distribution of the number of interaction domains (a), the total number (b) and the average span (c) of interaction domains detected in each cell, stratified by cell-types. Asterisks indicate the level of statistical significance of a pairwise paired Wilcoxon test against adipocytes; *** indicates adjusted *P*<0.05 and n.s. denotes non-significant. **d**, Scatter plot showing the short to long-range interaction ratio per cell against the number of interaction domains detected. Cells are colored by its snm3C-seq annotation. **e-f**, Scatter plots showing the aggregated cell-type level median number of UMIs detected in cells by snRNA-seq against the median number of interaction domains (e) and the ratio of short to long-range interaction contacts (f) detected in cells by snm3C-seq, colored similarly as in **d**. **g-h**, Bar plots showing the median distance (g) and the total number (h) of loop summits detected across the SAT cell-types (x-axis is ordered by the abundance in snm3C-seq). ASPC indicates adipose stem and progenitor cells.

**Extended Data Figure 7.**
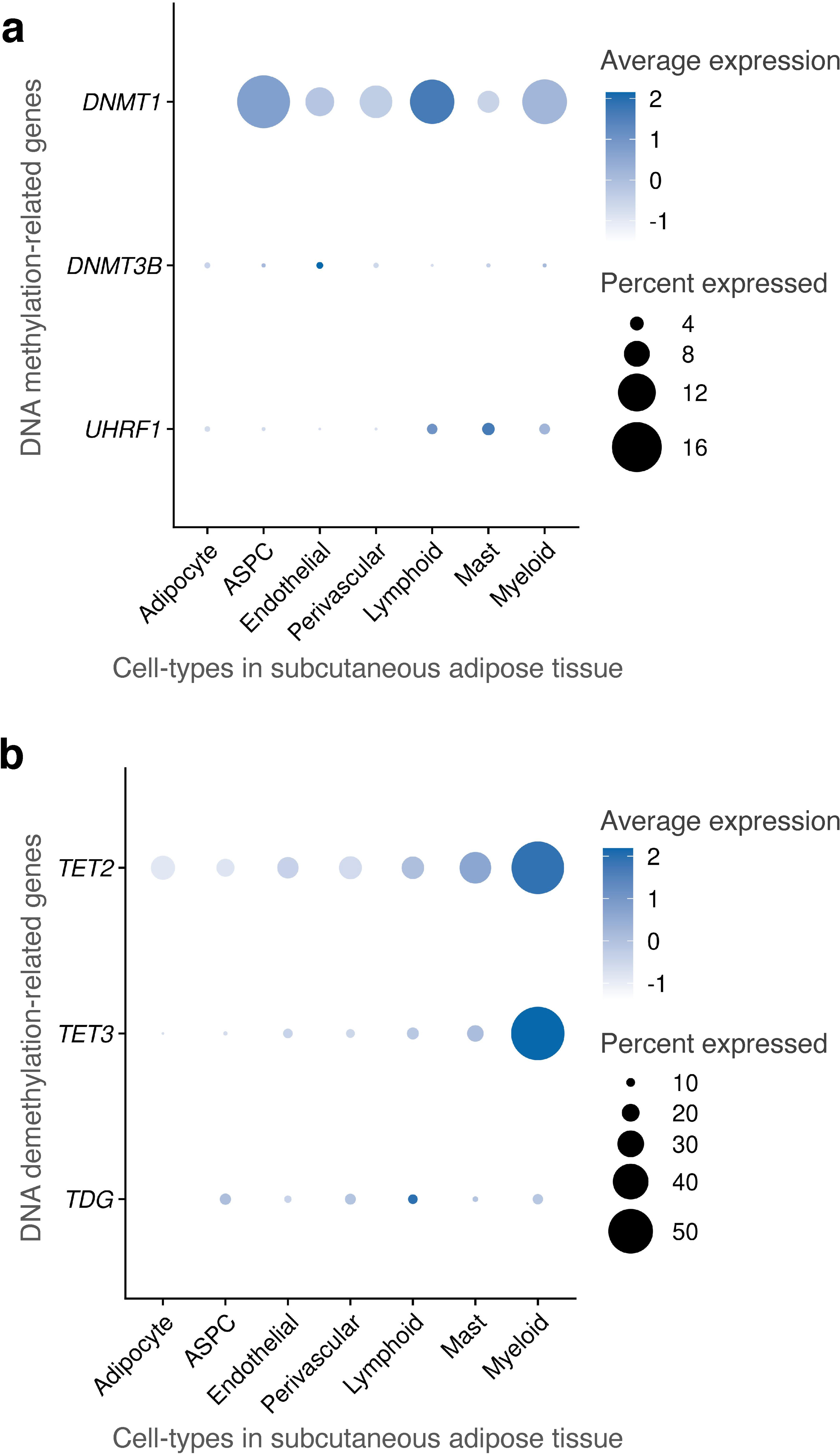
Mean gene expression of DNA methylation- and demethylation-related genes across cell-types in subcutaneous adipose tissue (SAT). **a-b**, Dot plot showing expression of (a) DNA methylation genes (*DNMT1, DNMT3B,* and *UHRF1*) and (b) DNA demethylation genes (*TET2*, *TET3*, and *TDG*) across subcutaneous adipose tissue (SAT) cell-types. The size of the dot represents the percentage of cells, in which a gene is expressed within a cell-type while the color represents the average expression of each gene across all cells within a cell-type (blue indicates a higher expression). ASPC indicates adipose stem and progenitor cells.

**Extended Data Figure 8.**
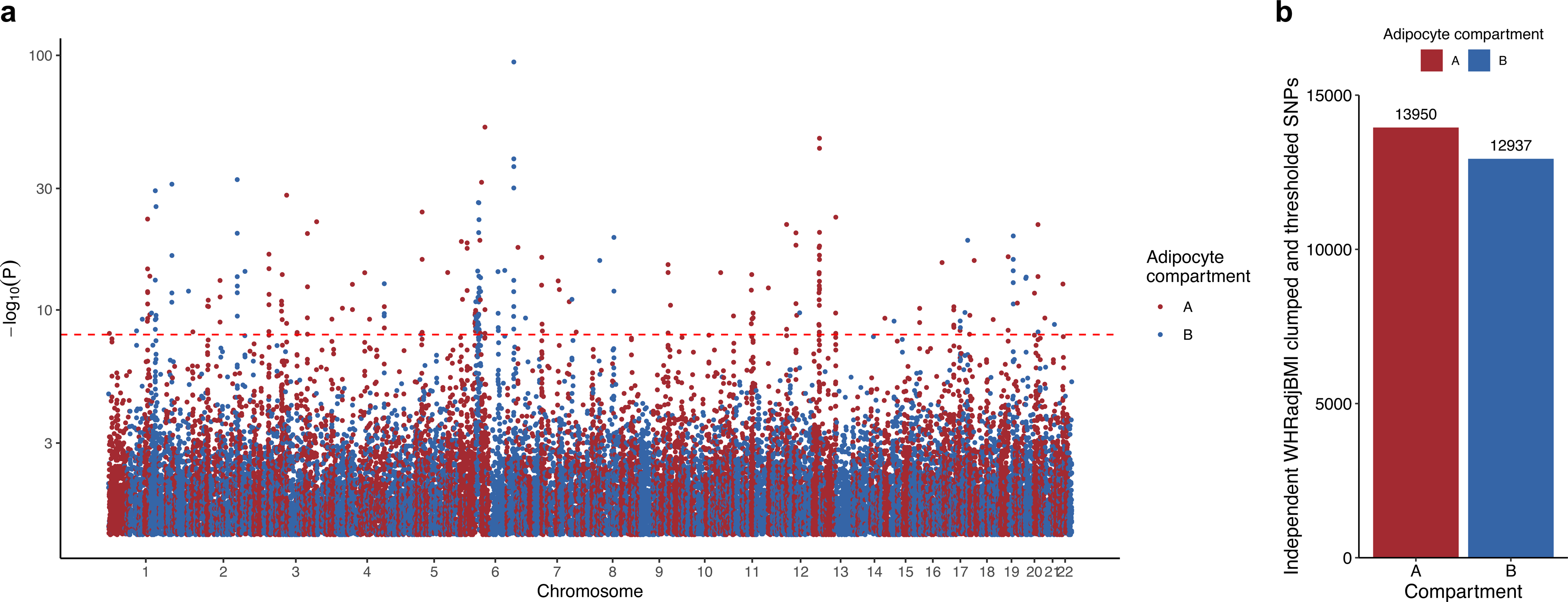
Abdominal obesity -associated variants are enriched for the adipocyte A compartment. a, The clumped and thresholded variants (r^2^<0.1, *P*<0.05) used for the adipocyte compartment PRSs for abdominal obesity (employing waist-hip-ratio adjusted for BMI (WHRadjBMI) as a proxy) are plotted by genomic position against the -log_10_*P* from the UK Biobank WHRadjBMI GWAS that we used for the WHRadjBMI PRS base (n=195,863 unrelated Europeans). SNPs landing in the adipocyte A compartment are colored blue, while SNPs landing in the adipocyte B compartment are colored black. **b,** Bar plot showing the number of independent (r^2^<0.1) WHRadjBMI-associated variants, passing nominal significance (*P*<0.05), from the WHRadjBMI GWAS, conducted in 195,863 individuals from the UK Biobank, grouped by the adipocyte compartment assignment.

