## Supplementary Table 3 for "Single-cell DNA methylome and 3D genome atlas of the human subcutaneous adipose tissue"

**Supplementary Table 3.** Numbers and proportions of differentially methylated regions (DMRs; hypo-methylated, n.s.*, and hyper-methylated) with their corresponding -log_10_*P*** in subcutaneous adipose tissue (SAT) cell-types.

| **SAT**  **cell-type** | **Hypo**  (%) | **Hypo**  **(-log_10_*P***) | **n.s.^*^**  (%) | **Hyper**  **(**%) | **Hyper**  **(-log_10_*P***) |
| --- | --- | --- | --- | --- | --- |
| Adipocyte | 396,758 (56.3%) | 129.0 | 208,464 (29.6%) | 99,841 (14.2%) | 0.0 |
| ASPC^#^ | 356,844 (50.6%) | 104.7 | 197,644 (28.0%) | 150,575 (21.4%) | 0.0 |
| Transition | 172,625 (24.5%) | 35.3 | 523,100 (74.2%) | 9,338 (1.3%) | 0.0 |
| Perivascular | 197,855 (28.1%) | 0.3 | 407,779 (57.8%) | 99,429 (14.1%) | 0.0 |
| Endothelial | 190,550 (27.0%) | 0.0 | 240,280 (34.1%) | 274,233 (38.9%) | 48.9 |
| Myeloid | 102,756 (14.6%) | 0.0 | 87,873 (12.5%) | 514,434 (73.0%) | 163.0 |
| Lymphoid | 93,984 (13.3%) | 0.0 | 417,090 (59.2%) | 193,989 (27.5%) | 6.7 |
| Mast | 64,026 (9.1%) | 0.0 | 451,258 (64.0%) | 189,779 (26.9%) | 19.5 |

*n.s. indicates not significantly hypo- or hyper-methylated.

***P* is calculated per hypo- and hyper-methylation state, jointly across all cell-types, using a one-tailed t-test adjusted for the number of cells in each cell-type (see Methods).

^#^ ASPC indicates adipose stem and progenitor cells.
