## Supplementary Table 5 for "Single-cell DNA methylome and 3D genome atlas of the human subcutaneous adipose tissue"

**Supplementary Table 5.** Numbers and proportions of the differential 100-kb bins for the top 5 most abundant cell-types in subcutaneous adipose tissue (SAT), identified by dcHiC using the aggregated pseudobulk level contact matrices, stratified by compartments (see Methods).

| SAT cell-type | A compartment | B compartment |
| --- | --- | --- |
| Adipocyte | 6,289 (54.35%) | 5,282 (45.65%) |
| ASPC^#^ | 6,327 (54.68%) | 5,244 (45.32%) |
| Endothelial | 6,127 (52.95%) | 5,444 (47.05%) |
| Perivascular | 6,342 (54.81%) | 5,229 (45.19%) |
| Myeloid | 6,161 (53.25%) | 5,410 (46.75%) |

^#^ ASPC indicates adipose stem and progenitor cells.
